# Uncovering the unexplored diversity of thioamidated ribosomal peptides in Actinobacteria using the RiPPER genome mining tool

**DOI:** 10.1101/494286

**Authors:** Javier Santos-Aberturas, Govind Chandra, Luca Frattaruolo, Rodney Lacret, Thu H. Pham, Natalia M. Vior, Tom H. Eyles, Andrew W. Truman

## Abstract

The rational discovery of new specialized metabolites by genome mining represents a very promising strategy in the quest for new bioactive molecules. Ribosomally synthesized and post-translationally modified peptides (RiPPs) are a major class of natural product that derive from genetically encoded precursor peptides. However, RiPP gene clusters are particularly refractory to reliable bioinformatic predictions due to the absence of a common biosynthetic feature across all pathways. Here, we describe RiPPER, a new tool for the family-independent identification of RiPP precursor peptides and apply this methodology to search for novel thioamidated RiPPs in Actinobacteria. Until now, thioamidation was believed to be a rare post-translational modification, which is catalyzed by a pair of proteins (YcaO and TfuA) in Archaea. In Actinobacteria, the thioviridamide-like molecules are a family of cytotoxic RiPPs that feature multiple thioamides, and it has been proposed that a YcaO-TfuA pair of proteins also catalyzes their formation. Potential biosynthetic gene clusters encoding YcaO and TfuA protein pairs are common in Actinobacteria but the chemical diversity generated by these pathways is almost completely unexplored. A RiPPER analysis reveals a highly diverse landscape of precursor peptides encoded in previously undescribed gene clusters that are predicted to make thioamidated RiPPs. To illustrate this strategy, we describe the first rational discovery of a new family of thioamidated natural products, the thiovarsolins from *Streptomyces varsoviensis*.

## INTRODUCTION

Microorganisms have provided humankind with a vast plethora of specialized metabolites with invaluable applications in medicine and agriculture.^1^ The advent of widespread genome sequencing has shown that the metabolic potential of bacteria had been substantially underestimated, as their genomes contain many more biosynthetic gene clusters (BGCs) than known compounds.^2,3^ Much of this enormous potential is either unexplored or undetectable under laboratory culture conditions, and is likely to include structurally novel bioactive specialized metabolites. Among the main classes of specialized metabolites produced by microorganisms, the ribosomally synthesized and post-translationally modified peptides^4^ (RiPPs) may harbor the largest amount of unexplored structural diversity. This is due to the inherent difficulties related to the *in silico* prediction of their BGCs, as RiPP biosynthetic pathways lack any kind of universally shared feature apart from the existence of a pathway-specific precursor peptide.

RiPP BGCs can be identified by the co-occurrence of specific RiPP tailoring enzymes (RTEs) alongside a precursor peptide that contains sequence motifs that are characteristic of a given RiPP family. This makes it relatively simple to identify further examples of known RiPP families,^5,6^ but the identification of currently undiscovered RiPP families remains a significant unsolved problem. Unlike specialized metabolites such as polyketides, non-ribosomal peptides and terpenes, there are no genetic features that are common to all RiPP BGCs to aid in their identification. Furthermore, genes encoding precursor peptides are often missed during genome annotation due to their small size, yet the reliable prediction of precursor peptides constitutes a crucial task, as this starting scaffold is essential for RiPP structural prediction. Numerous analyses of specific RiPP classes signal the existence of a wide array of uncharacterized RiPP families,^7-9^ but currently available prediction tools still rely on precursor peptide features that are associated with known RiPP families, thereby limiting the discovery of new RiPP families.^10-14^

YcaO domain proteins are a widespread superfamily of enzymes with an intriguing catalytic potential in RiPP biosynthesis.^15^ These were originally shown to be responsible for the introduction of oxazoline and thiazoline heterocycles in the PP backbone of microcins,^16^ and were very recently demonstrated to catalyze the formation of the macroamidine ring of bottromycin.^17-19^ YcaO proteins act as cyclodehydratases, activating the amide bond substrate by nucleophilic attack, which is followed by ATP-driven O-phosphorylation of the hemiorthoamide intermediate and subsequent elimination of phosphate. In most azoline-containing RiPPs, this catalytic activity requires a partner protein (E1-like or Ocin-ThiF-like proteins that are clustered with or fused to the YcaO domain), which acts as a docking element to bring the precursor peptide to the active site of the cyclodehydratase. YcaO proteins can also act as standalone proteins, as in bottromycin biosynthesis,^17-19^ and many YcaO proteins are encoded in genomes without E1-like or Ocin-ThiF-like partner proteins,^9,15^ including in the BGCs of thioviridamide-like molecules.^6,20-24^

Thioviridamide and related compounds are cytotoxic RiPPs that contain multiple thioamide groups (Figure 1), but no azole or macroamidine rings. Thioamides are rare in nature^25-31^ and it has been hypothesized that YcaO proteins could be responsible for this rare amide bond modification in thioviridamide biosynthesis, potentially in cooperation with TfuA domain proteins^15^ (Figure 1). This protein pair has been identified elsewhere in nature, including in archaea, where they are involved in the ATP-dependent thioamidation of a glycine residue of methyl-coenzyme M reductase.^32,33^ We therefore hypothesized that the identification of *tfuA*-like genes could be employed as a rational criterion for the identification of BGCs responsible for the production of novel thioamidated RiPPs in bacteria.

**Figure 1.**
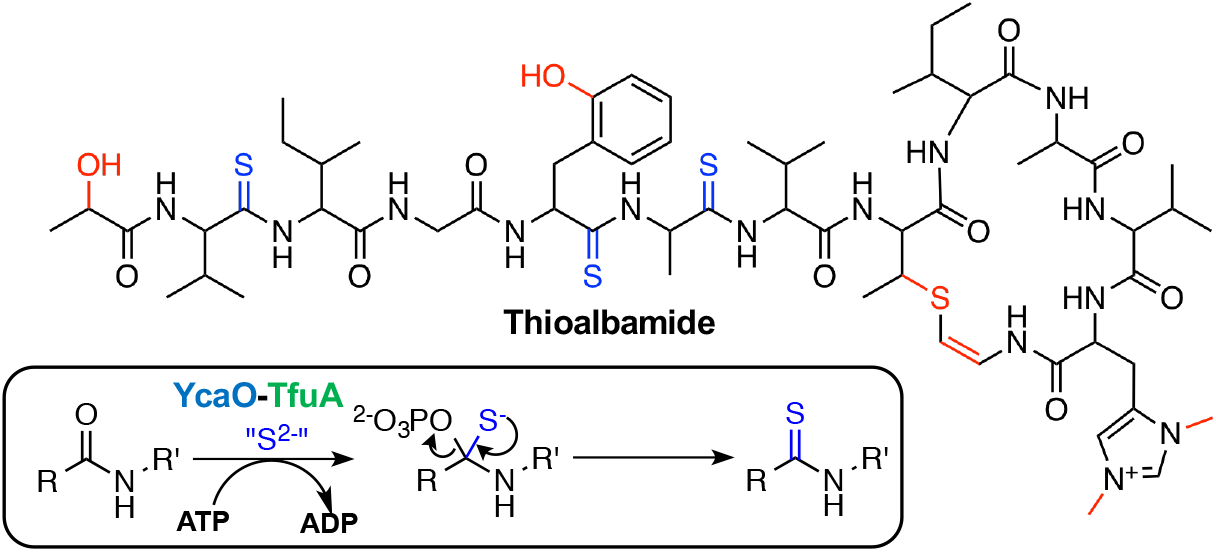
An example of a thioviridamide-like molecule, thioalbamide, and inset, a proposed biochemical route to thioamides. Thioamides are highlighted in blue and other post-translational modifications are colored red.

An exploration of the diversity of *tfuA*-containing BGCs required methodology to identify precursor peptides that have no homology to known precursor peptides. Here, we report RiPPER (RiPP Precursor Peptide Enhanced Recognition), a method for the identification of precursor peptides that requires no information about RiPP structural class (available at https://github.com/streptomyces/ripper). This evaluates regions surrounding any putative RTE for short open reading frames (ORFs) based on the likelihood that these are truly peptide-coding genes. Peptide similarity networking is then used to identify putative RiPP families. We apply this methodology to identify RiPP BGCs encoding TfuA proteins in Actinobacteria, which reveals a highly diverse landscape of BGC families that are predicted to make thioamidated RiPPs. This analysis informed the discovery of the thioamidated thiovarsolins from *Streptomyces varsoviensis*, which are predicted to belong to a wider family of related thioamidated RiPPs and represents the first rational discovery of a new family of thioamidated compounds from nature.

## RESULTS AND DISCUSSION

### Development of a family-independent RiPP genome mining tool

Within a given RiPP family, all BGCs usually encode at least one tailoring enzyme and one precursor peptide that each feature domains conserved across the RiPP family.^4^ This has led to the development of genome mining methodology that can identify these well-characterized RiPP families with high accuracy.^11-13^ However, there is a growing number of widespread RiPP BGCs with little or no homology to known RiPP BGCs.^7,34^ Theoretically, backbone modification like thioamidation or epimerization^35^ can occur on any residue. In addition, well-characterized RiPP tailoring enzymes can be associated with unusual precursor peptides that lack homology to known RiPP classes.^9^ We therefore sought to develop a method to identify likely precursor peptides that was independent of PP sequence and could be applicable for any RiPP family. The starting point for this method was to employ the functionality of RODEO^13, 14^ to identify genomic regions associated with a series of putative RTEs. RODEO uses a mixture of heuristic scoring and support vector machine classification to identify precursor peptides for lasso peptides^13^ and thiopeptides,^14^ but does not accurately identify other precursor peptides, whose sequences are highly variable and are often not annotated in genomes.

To enable the sequence independent discovery of precursor peptides, we sought to identify short ORFs that possess similar genetic features as other genes in a given gene cluster, including ribosome binding sites, codon usage and GC content. Prodigal (PROkaryotic DYnamic programming Gene-finding ALgorithm) uses these criteria to identify bacterial ORFs.^36^ Therefore, following RODEO retrieval of nucleotide data, we implemented a modified form of this algorithm to specifically search for ORFs that encode for peptides of between 20 and 120 amino acids within apparently non-coding regions near to a predicted RTE (Figure 2A). Given the prevalence of characterized precursor peptides that are encoded on the same strand as a tailoring gene, a same strand score is added (custom parameter; default = 5). A modified GenBank file is generated by RiPPER that annotates these putative short ORFs within the putative BGC (Figure S1), and these are ranked alongside annotated short genes based on their Prodigal score. RiPPER then retrieves the top three scoring ORFs within ±8 kb of the RTE, plus any additional high scoring ORFs over a specified score threshold that represent probable genes. These are then assessed for Pfam domains^37^ and data associated with each peptide is tabulated for further processing.

**Figure 2.**
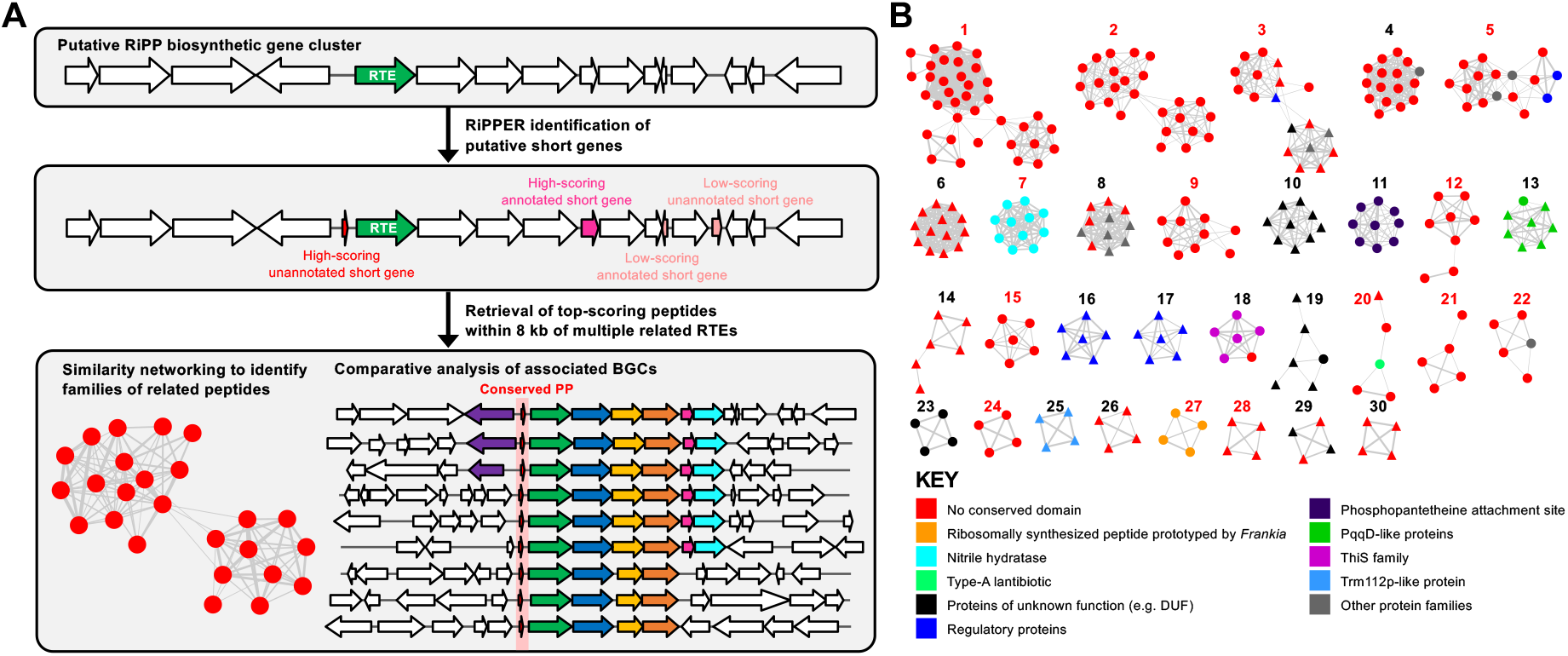
RiPPER identification of putative precursor peptides. (A) Schematic of RiPPER workflow where a cluster is identified based on a putative RiPP tailoring enzyme (RTE). (B) The 30 largest peptide similarity networks identified using RiPPER for peptides associated with *tfuA-like* genes in Actinobacteria. Red numbers indicate networks predicted to comprise of authentic precursor peptides (see Table S1 and Figures S6-S19) and triangular nodes indicate peptides encoded on the opposite strand to the RTE gene.

To validate this approach, we used RTE accession numbers that had previously been used to identify lasso peptide (RODEO^13^), microviridin^38^ and thiopeptide (RODEO^14^) gene clusters. In each case, class-specific rules had been used to identify associated precursor peptides. These RiPP classes are well-suited to method validation as they have diverse gene cluster features and precursor peptide sequences, and span multiple bacterial taxa. In addition, the genes encoding these small peptides are often not annotated in genome sequences.^13^ We therefore used RiPPER with the same protein accessions as those previous studies to retrieve BGCs and associated precursor peptides. Comparison of the RiPPER outputs with these studies revealed that lasso peptide and microviridin precursor identification was highly reliable. 1056 out of 1122 (94.1%) and 279 out of 288 (96.7%) peptides identified by those prior mining studies were identified by RiPPER (Table 1, Supplementary Datasets 1-2).

**Table 1.** Comparison of RiPPER with prior studies on the identification of RiPP precursor peptides.

| RiPP class<sup>a</sup> | No. of RTEs used in RiPPER search | Unbiased RiPPER search |  | RiPPER including HMM search |  | Network 1 data from RiPPER analysis |  |  |
| --- | --- | --- | --- | --- | --- | --- | --- | --- |
|  |  | Total peptides retrieved | Match with prior data<sup>b</sup> | Total peptides retrieved | Match with prior data<sup>b</sup> | Total peptides in network | Match with prior data<sup>b</sup> | Additional HMM hits |
| Lasso peptides | 1198 | 4503 | 1056/1122 (94.1%) | 4558 | 1064/1122 (94.8%) | 1211<sup>c</sup> | 934/1122 (83.2%) | 125 |
| Microviridins | 159 | 586 | 270/280 (96.4%) | 596 | 270/280 (96.4%) | 270 | 269/280 (96.1%) | 1 |
| Thiopeptides | 486 | 1526 | 439/591 (74.3%) | 1675 | 550/591 (93.1%) | 690 | 543/591 (91.9%) | 75 |

<sup>a</sup> Data obtained for lasso peptides from ref. 13, microviridins from ref. 38 and thiopeptides from ref. 14.
<sup>b</sup> These numbers are sometimes greater than the number of RTEs used in the RiPPER search due to the identification of multiple precursor peptides per BGC.
<sup>c</sup> Proteins with PqqD domains removed.

In contrast, RiPPER only retrieved 439 of the 591 (74.3%) thiopeptide precursors previously identified (Table 1, Supplementary Dataset 3). This was possibly due to the comparatively large size of thiopeptide BGCs, which meant that the ±8 kb search window was not suited to a subset of these BGCs. Widening the unbiased search reduced specificity of the retrieval, so an additional targeted search step was introduced. All short peptides across the entire gene cluster region (default = 35 kb) that were not retrieved by the first search were analyzed for precursor peptide domains using hidden Markov models (HMMs) recently built by Haft *et al*.^39^ Any peptides containing a domain were therefore also retrieved. This provided a minor improvement to RiPPER retrieval of lasso precursor peptides but significantly improved thiopeptide precursor peptide retrieval to 543 out of 591 (91.9%) peptides identified by RODEO.^14^

This data demonstrated that the RiPPER methodology was applicable to multiple diverse classes of RiPP, but the unbiased nature of retrieval meant that only between a half and a quarter (depending on RiPP class) of total retrieved peptides were likely to be precursor peptides (Table 1). We therefore generated peptide similarity networks^40^ using peptides retrieved from each RiPPER analysis, where peptides with at least 40% identity were connected to each other. Despite the large sequence variance within each RiPP class, this was highly effective at filtering the peptides into networks of likely precursor peptides. For each RiPPER analysis, the largest network (“network 1”) contained the majority of precursor peptides identified by previous studies (Table 1, Figures S2-S4). Unexpectedly, network 1 of the lasso peptide dataset also contained PqqD domain proteins, a conserved feature of lasso peptide pathways that function as RiPP precursor peptide recognition elements (RREs).^41,42^ These peptides could be easily filtered by Pfam analysis, as would a higher identity cut-off. In addition, network 2 comprises of 56 *Burkholderia* peptides that are precursors to capistruin lasso peptides (all identified by RODEO). Notably, for each RiPPER analysis, network 1 contained peptides with the expected precursor peptide domain that were not retrieved by either RODEO^13,14^ or the bespoke microviridin analysis.^38^ In total, this provided over 200 new candidate precursor peptides (Table 1), as well as additional networked peptides with no known domains that could feasibly be authentic precursor peptides. The ability of RiPPER to correctly identify a comparable number of precursor peptides to prior targeted methods demonstrates that the combination of rational ORF identification and scoring, Pfam analysis, and peptide similarity networking can identify RiPP precursor peptides with a high degree of accuracy and coverage without any prior knowledge of the RiPP class.

### Identification of thioamidated RiPP BGCs using RiPPER

As a backbone modification, thioamidation potentially has no requirement for specific amino acid side chains, which means that there may be no conserved sequence motifs within precursor peptide substrates. To guide our identification of thioamidated RiPP BGCs, we identified a curated set of 229 TfuA-like proteins in Actinobacteria whose putative BGCs were retrieved using RiPPER, which showed that each TfuA protein was encoded alongside a YcaO protein but their associated gene clusters could be highly variable. RiPPER retrieved 743 peptides (Supplementary Dataset 4) and peptide similarity networking (40% identity cut-off) yielded 74 distinct networks of peptides, where 30 of these networks featured four or more peptides (Figure 2B, Figure S5, Supplementary Table 1). MultiGeneBlast^43^ was then employed to compare the BGCs corresponding to each network.

As an initial proof of concept, this correctly grouped all thioviridamide-like precursor peptides into a single network (Figure 3A). Surprisingly, these precursor peptides were connected with four additional peptides encoded in putative BGCs that are extremely different to thioviridamide-like BGCs; three of these peptides were not previously annotated as genes. These peptides feature extensive sequence similarities with the thioviridamide-like precursor peptides (Figure S6), but the BGCs themselves are extremely different, where the only common features with the thioviridamide-like BGCs are the YcaO, TfuA and precursor peptide genes (Figure 3B). More generally, peptide networking guided the identification of a wide variety of probable tfuA-containing RiPP BGCs (Figures S6-S19). For example, many mycobacteria encode a YcaO-TfuA protein pair, and the largest network of putative precursor peptides is associated with this mycobacterial BGC (Figure 2B, Network 1) where they are usually encoded near a Type III polyketide synthase (PKS) and a sulfotransferase (Figure S7). Network 2 consists of 25 related *Streptomyces* peptides that possess high Prodigal scores and are encoded at the start of a conserved biosynthetic operon (Figure S8). This is a strong candidate as an authentic RiPP BGC family, yet only 6 of these 25 short peptides were originally annotated.

**Figure 3.**
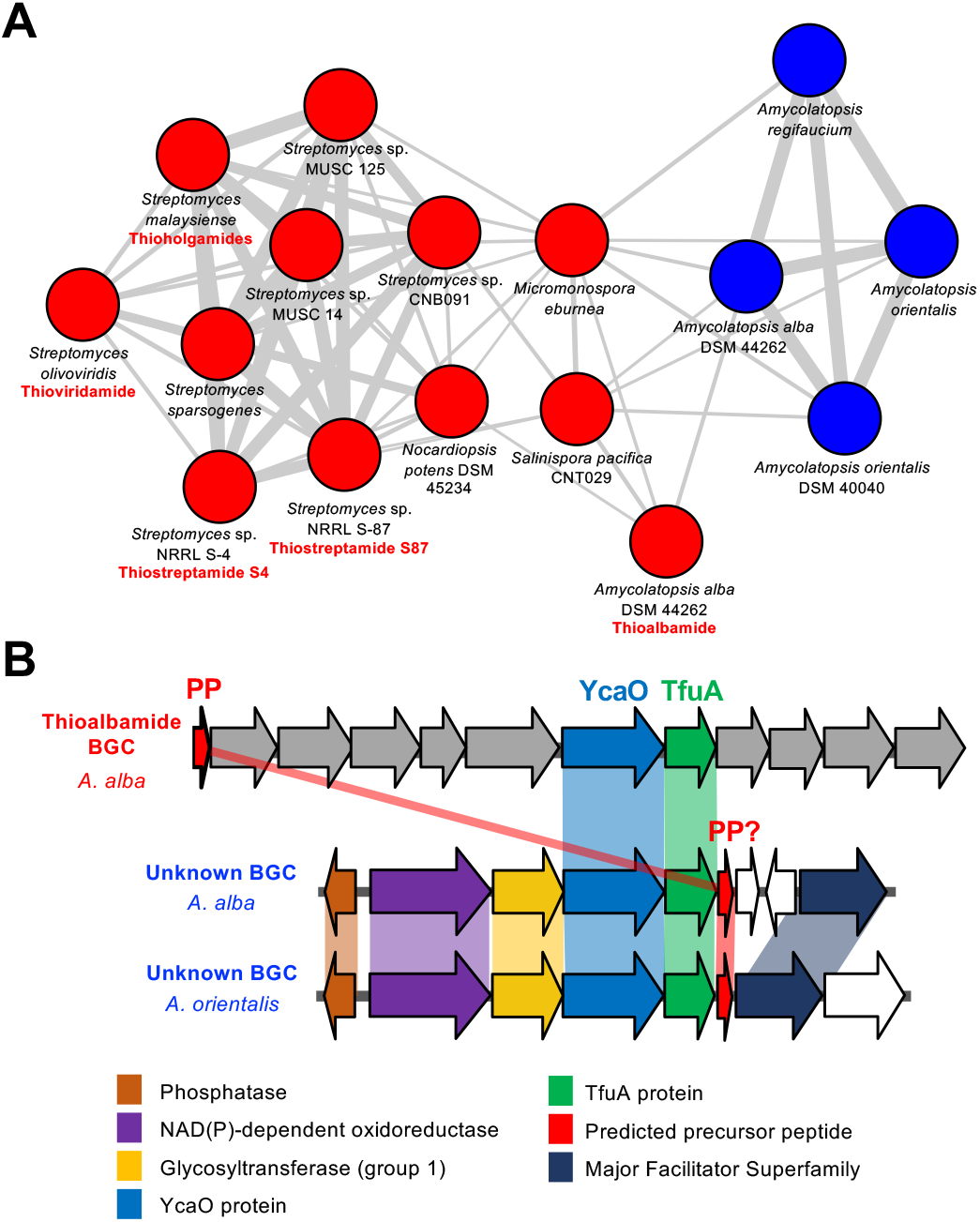
Thioviridamide-like precursor peptides. (A) The precursor peptide network that includes both thioviridamide-like precursor peptides (red nodes) and a related but uncharacterized family of precursor peptides from BGCs that are highly different to thioviridamide-like BGCs (blue nodes). Characterized compounds are listed with their respective nodess. (B) Comparative analysis of thioviridamide-like and non-thioviridamide-like BGCs from this network where related genes share the same color. See Figure S6 for full BGC details.

### Thioamidated RiPPs are a largely unexplored area of the natural products landscape

To investigate whether BGC families correlate with the evolutionary relationships of the TfuA proteins, a maximum likelihood tree was constructed from standalone TfuA domain proteins and the peptide networks were mapped to this tree (Figure 4, Supplementary Dataset 5). This showed strong correlations between TfuA phylogeny and precursor peptide similarity. Despite the significant differences between their gene clusters, the thioviridamide-like and non-thioviridamide-like peptides of Network 5 are all associated with closely related TfuA proteins. Unsurprisingly, some TfuA domain proteins are associated with multiple peptide networks due to the abundance of small peptides that are unlikely to be precursor peptides, such as regulatory proteins and RREs.^42^ For example, almost all peptides from Networks 9, 11 and 18 are associated with the same set of TfuA domain proteins, but Pfam analysis indicates that Networks 11 and 18 consist of acyl carrier proteins and ThiS-like proteins,^44^ respectively.

**Figure 4.**
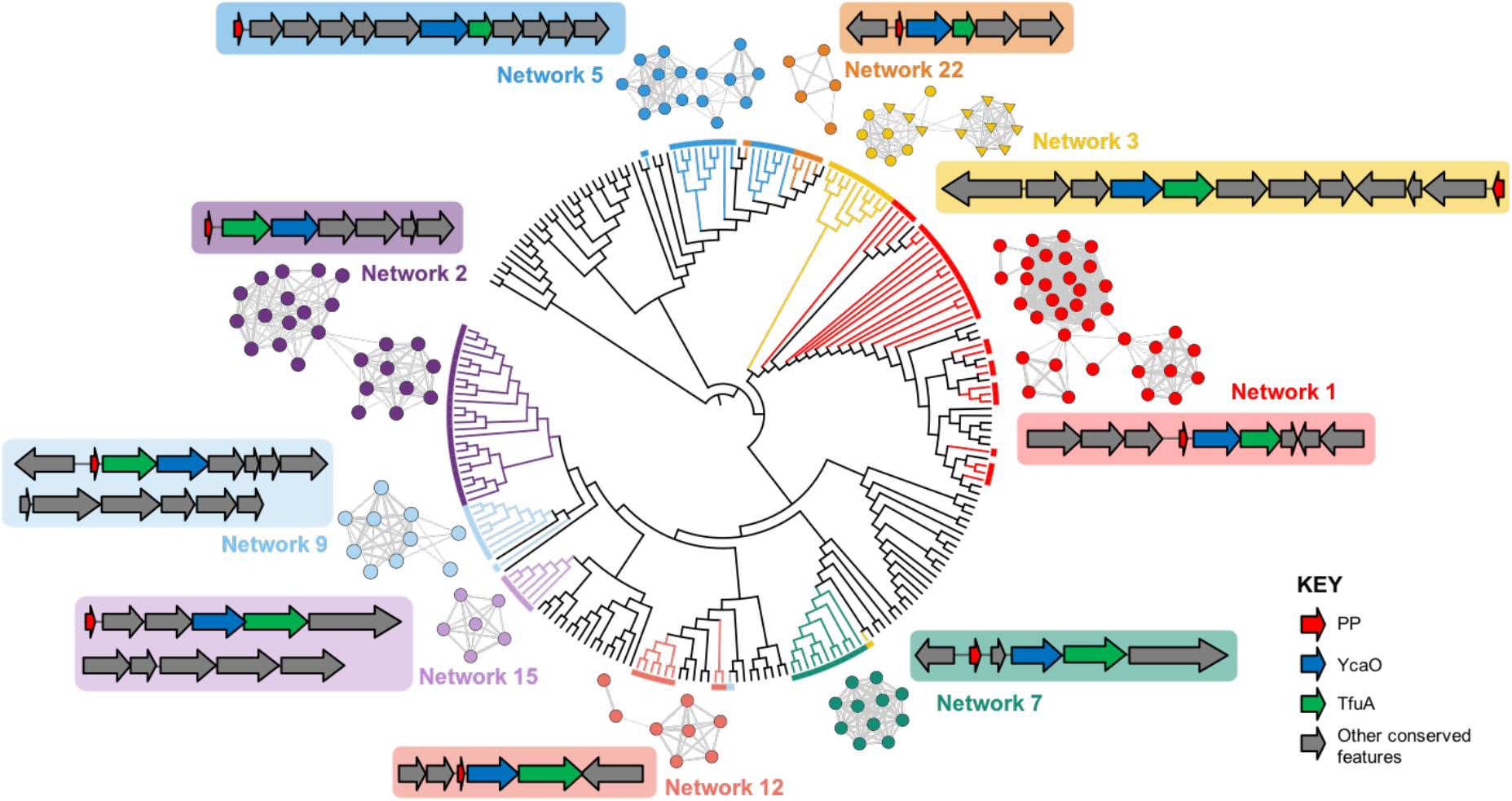
Examples of putative RiPP BGCs and associated TfuA phylogeny. A maximum likelihood tree (branch lengths removed) of TfuA-like proteins is color-coded to indicate the relationship between TfuA-like proteins and the associated networks of putative precursor peptides. Representative BGCs are also shown, where grey genes indicate genetic features that are conserved across multiple BGCs within that family. Fully annotated BGCs are shown in Figures S6-S19.

Therefore, the Network 9 peptides, which are encoded at the beginning of each BGC and feature no conserved domains, are likely precursor peptides for this BGC family (Figure 4). In contrast, Pfam analysis indicated that all precursor peptides in Network 7 feature nitrile hydratase domains, which is a common feature amongst precursor peptides across diverse RiPP families.^8,45^ In total, at least 15 distinct predicted RiPP families were predicted from the top 30 peptide networks (Supplementary Dataset 4, Table S1, Figures S6-S19), while many smaller networks and singletons are also likely to be authentic precursor peptides, based on their Prodigal scores and positions within BGCs. A comparative analysis with the source GenBank entries indicated that over half of the peptides encoded in these BGCs were not previously annotated (Supplementary Dataset 4); on average, unannotated peptides identified by RiPPER were significantly shorter than annotated peptides (Figure S20).

### Characterization of a novel family of TfuA-YcaO BGCs

To determine whether the newly identified YcaO-TfuA BGCs actually produce thioamidated RiPPs, we focused on Network 22 (Figure 5A), a group of five orphan BGCs with multiple unusual features (Figure 5B). Most notably, the predicted precursor peptides feature a series of imperfect repeats that could reflect a repeating core peptide (Figure 5C), where the family varies from a non-repeating precursor peptide (*Asanoa ishikariensis*) to five repeats (*Streptomyces varsoviensis*). In addition, the *Nocardiopsis* and *Streptomyces* BGCs encode two additional conserved proteins, an amidinotransferase (AmT) and an ATP-grasp ligase, which are homologous to proteins in the pheganomycin pathway,^46^ and are adjacent to genes encoding non-ribosomal peptide synthetases (NRPSs) or PKSs (Figure 5B). Efforts to genetically manipulate *S. varsoviensis* and *Nocardiopsis baichengensis* were unsuccessful and we were unsure of the gene cluster boundaries, so transformation-associated recombination (TAR) cloning^47,48^ was employed to capture a 31.7 kb DNA fragment comprising 25 genes (Table S2) centered around the *ycaO-tfuA* core of the *S. varsoviensis* BGC. Two independent positive TAR clones were conjugated into three different host strains: *Streptomyces lividans* TK24 and *Streptomyces coelicolor* M1146 and M1152^49^ and the resulting TARvar exconjugants were fermented in a variety of media. Liquid chromatography-mass spectrometry (LC-MS) analysis revealed two major compounds (*m/z* 399.18 and *m/z* 401.20), and two minor compounds (*m/z* 385.16 and *m/z* 387.18) not present in the negative control strains (Figure 5D). Small amounts of these compounds could be detected when *S. varsoviensis* was fermented for 10 days (Figure 6, Figure S21).

**Figure 5.**
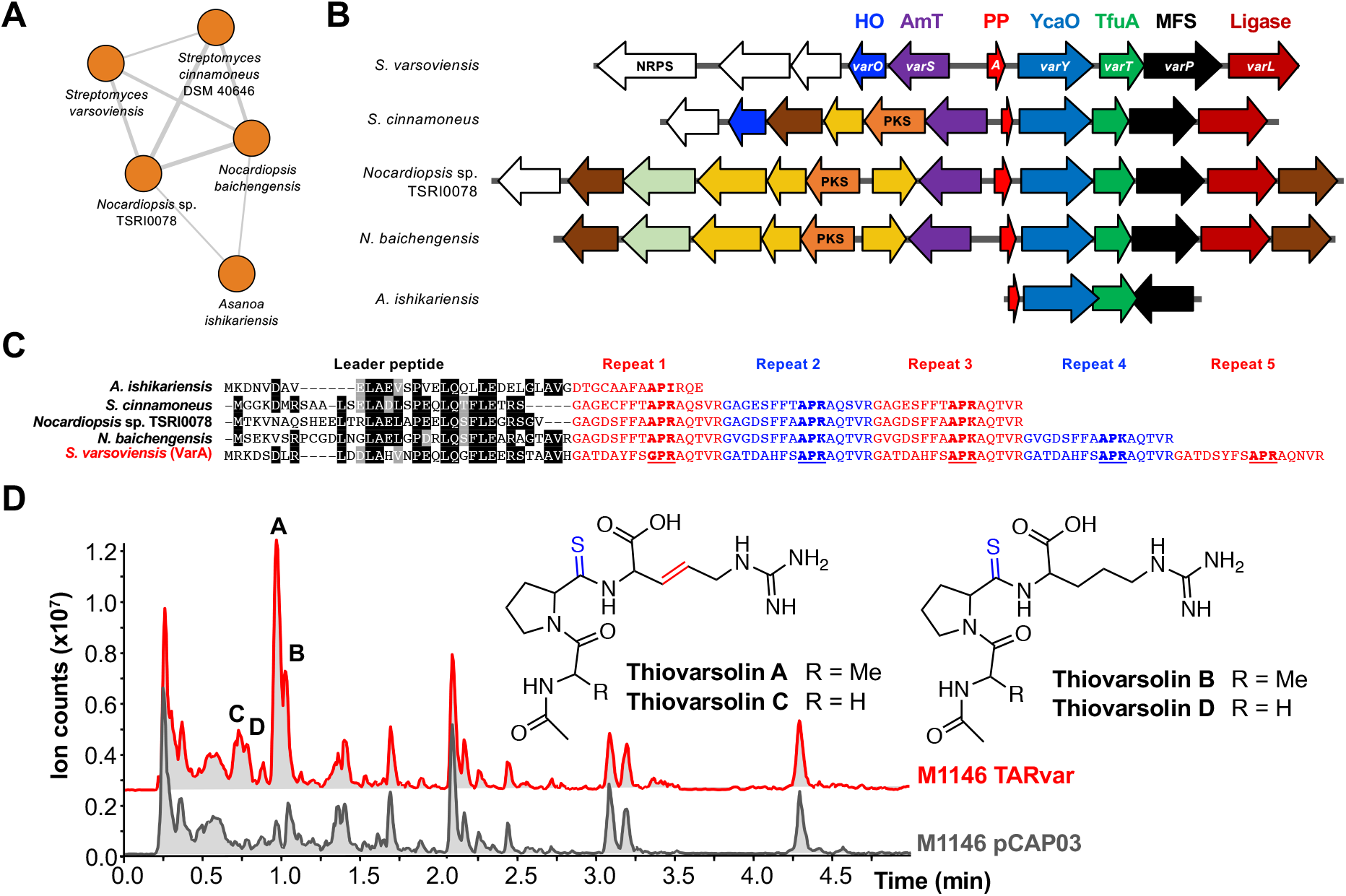
Identification of the thiovarsolin family of RiPPs. (A) The associated precursor peptide network. (B) BGCs associated with each precursor peptide. The protein product of each *var* gene is listed at the top (HO = heme oxygenase; AmT = amidinotransferase; MFS = major facilitator superfamily) and genes common to multiple BGCs are color-coded by the predicted function of the protein product (see Figure S16 for full details). (C) Putative repeating precursor peptides identified by similarity networking. The predicted leader peptide is aligned, while the repeat regions are highlighted. Underlined text indicates the partially conserved core peptide that the thiovarsolins derive from, and bold text indicates equivalent residues in the other precursor peptides. (D) Analysis of thiovarsolin production by *S. coelicolor* M1146 TARvar, which contains a 31.7 kb DNA fragment centered on the *S. varsoviensis* BGC. Base peak chromatograms of crude extracts of *S. coelicolor* M1146 TARvar and an empty vector negative control (pCAP03) are shown, with peaks corresponding to thiovarsolins A-D indicated. Thioamidation and dehydrogenation post-translational modifications are highlighted on the thiovarsolin structures.

**Figure 6.**
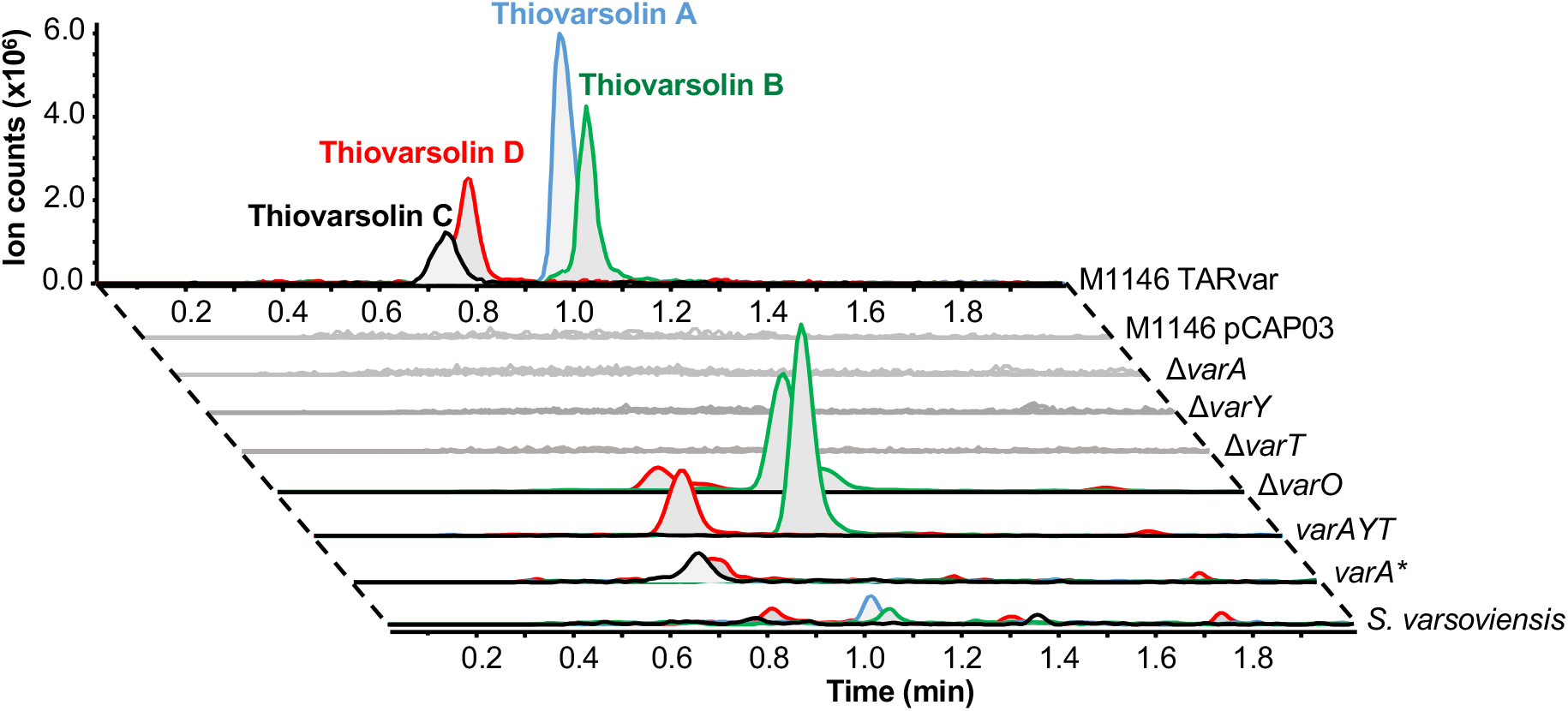
Mutational analysis of thiovarsolin biosynthesis. Extracted ion chromatograms (EICs) are shown for each thiovarsolin (A = *m/z* 399.18, B = *m/z* 401.20, C = *m/z* 385.16, D = *m/z* 387.18). M1146 pCAP03 indicates the empty plasmid control, while each ∆*var* mutation was made in the TARvar construct and expressed in *S. coelicolor* M1146. See text and Figure S35 for details of *varA*\*.

To associate the production of these new compounds to the cloned DNA fragment, PCR-targeting mutagenesis^50^ was employed to generate a series of deletion mutants on the putative BGC. A progressive trimming process determined that a cluster of seven genes that are conserved across the *Nocardiopsis* and *Streptomyces* BGCs was sufficient for compound production: *varA* (encoding the predicted repeating precursor peptide), *varY* (the YcaO protein), *varT* (the TfuA protein), *varO* (a heme oxygenase-like protein^51^), *varL* (an ATP-grasp ligase), *varP* (a major facilitator superfamily transporter) and *varS* (an amidinotransferase). The deletion of *varA*, *varY* and *varT* completely abolished the production of the four new compounds, while the ∆*varO* mutant produced only *m/z* 401.20 and *m/z* 387.18, suggesting that VarO may function as a dehydrogenase (Figure 6). Deletion of *varL*, *varP* and *varS* did not affect production, despite their conservation in related BGCs (Figure 5B). ∆*varY*, ∆*varT* and ∆*varO* mutants were successfully complemented by expressing these genes under the control of the *ermE*\* promoter, whereas complementation of ∆*varA* required its native promoter. As expected, expression of a 3.7 kb DNA fragment including only *varA*, *varY* and *varT* in *S*. *coelicolor* M1146 led to the production of *m/z* 401.20 and *m/z* 387.18 (Figure 6, *varAYT*). Collectively, this data show that *varAYTO* are the only genes required for the biosynthesis of this new group of RiPPs, thiovarsolins A-D (observed *m/z* 399.1818, 401.1968, 385.1652 and 387.1808, respectively, Table S5).

### The thiovarsolins are thioamidated peptides that derive from the repetitive core of the precursor peptide

The structures of thiovarsolins A and B were determined by NMR (^1^H, ^13^C, COSY, HSQC and HMBC; Figures S22-S33, Table S6) following large scale fermentation and purification of each compound. This analysis showed that thiovarsolins A and B are *N*-acetylated APR tripeptides in which the amide bond between Pro and Arg is substituted by a thioamide (δ_C_ 200 ppm) (Figure 5D). This was supported by accurate mass data (Table S5) and an absorbance maximum at ~270 nm for both molecules, which is characteristic of a thioamide group.^52^ Additionally, a trans double bond is present between Cβ and Cγ of the arginine side chain in thiovarsolin A. This peptide backbone is fully compatible with an APR sequence within the repeats of VarA (Figure 5C). The name thiovarsolin corresponds to linear thioamidated peptides made by *S. varsoviensis*.

Tandem MS (MS^2^) analysis of the thiovarsolins (Figure S34) revealed a clear structural relationship between thiovarsolins A (*m/z* 399.18) and C (*m/z* 385.16), as well as between thiovarsolins B (*m/z* 401.20) and D (*m/z* 387.18), which suggested that each 14 Da mass difference could be due to one methyl group. Interestingly, the first repetition of the putative modular core peptide features a GPR motif instead of APR, which could potentially explain this 14 Da mass difference, as well as their observed abundances in relation to thiovarsolins A and B. To test this hypothesis, a mutated version of *varA* was constructed (*varA*\*, Figure S35) in which the Ala residue in each repeat was substituted by Gly. This was expressed in M1146 TARvar ∆*varA* using a pGP9-based expression plasmid.^53^ The resulting strain was only able to produce thiovarsolins C and D (Figure 6, *varA*\*), confirming that these two minor compounds derive from a GPR core peptide. Such an extensively repeating precursor peptide is rare, but is comparable to the variable repeats found in precursor peptides for some cyanobactins^54^ and the fungal RiPP phomopsin.^55^

Our genetic and chemical analysis of the *var* BGC strongly suggests that the YcaO (VarY) and TfuA (VarT) proteins cooperate to introduce a thioamide bond. Given the absence of a specific protease in the gene cluster, it is plausible that endogenous peptidases are responsible for the liberation of the non-degradable thioamidated APR and GPR tripeptides, which later undergo an *N*- terminal acetylation catalyzed by an endogenous *N*-acetyltransferase, as previously reported for other metabolites containing primary amines.^56,57^ The timing of VarO-catalyzed dehydrogenation is unclear and could happen directly on the precursor peptide or after proteolysis. Small amounts of thiovarsolins A and B are produced by *S. varsoviensis*, but the lack of a function for *varS* and *varL* suggests that the described thiovarsolins might not be the final products of these pathways. However, no further thiovarsolin-related metabolites could be detected In either *S. varsoviensis* or *S. coelicolor* M1146 TARvar when analyzed by comparative metabolomics and by assessment of MS^2^ data for losses of H_2_S (*m/z* 33.99), which is a fragmentation profile that is characteristic of thioamides.^6^

## CONCLUSION

The discovery of the thiovarsolins supports the existence of an unexplored array of thioamidated RiPPs in Actinobacteria. The discovery that a minimal gene set of *varA* (precursor peptide), *varY* (YcaO protein) and *varT* (TfuA protein) is sufficient for the biosynthesis of thiovarsolin B (Figure 6) provides strong evidence that the YcaO-TfuA protein pair catalyze peptide thioamidation in bacteria, which is supported by a parallel study by Mitchell and colleagues on thiopeptide thioamidation.^14^ It was previously determined that a distantly related pair of homologs catalyze thioamidation of methyl-coenzyme M reductase in archaea.^32,33^ The relatively simple thiovarsolin pathway therefore represents a promising system for future biochemical studies of this reaction in the context of RiPP biosynthesis. Unexpectedly, genes conserved across multiple homologous var-like pathways (*varS*, *varP* and *varL*, Figure 5B) were not required for thiovarsolin biosynthesis. Along with N-terminal acetylation, this suggests that the identified thiovarsolins may be shunt products, although the production of thiovarsolins by *S. varsoviensis* indicates that they are made naturally, so production is not simply a consequence of heterologous pathway expression. The introduction of a double bond in the arginine residue side chain of the thiovarsolins by VarO would represent new RiPP biochemistry, as heme oxygenases have never been associated with RiPP biosynthesis. This shows that the breadth and diversity of RiPP post-translational modifications is still expanding, which has also been highlighted by recent discoveries of radical SAM enzyme-catalyzed epimerization,^45^ decarboxylation^58^ and β-amino acid formation^59^ in RiPP pathways.

RiPPER is a flexible prediction tool that can be applied to any class of predicted RiPP tailoring enzyme to aid in the discovery of this metabolic dark matter. This more general approach complements existing genome-mining tools such as BAGEL,^10^ RODEO,^13,14^ PRISM^60^ and antiSMASH,^12^ which all provide in-depth analyses and product predictions for established RiPP families. The *de novo* identification of precursors to lasso peptides, microviridins and thiopeptides highlights the scope of RiPPER, which was achieved without any specific rules for these RiPP families. The methodology proved to be highly adept at identifying previously overlooked precursor peptide genes, and the method parameters can be easily adapted based on prior knowledge of a given RiPP family (min/max gene length, max distance from RTE, same strand score and peptide score threshold, for example). In our TfuA analysis, peptide networking proved to be a highly effective method to prioritize related precursor peptides and their associated BGCs for further analysis, where it highlighted the existence of likely RiPP families as opposed to the coincidental presence of a small ORF near a putative BGC. The diversity of TfuA-associated precursor peptides identified in Actinobacteria highlights the utility of an unbiased precursor peptide identification tool and provides the basis for investigating the breadth of this RiPP family. It will be fascinating to determine both the structure and function of these cryptic metabolites.

## Supporting information

## AVAILABILITY

RiPPER is available at:

https://github.com/streptomyces/ripper and https://hub.docker.com/r/streptomyces/ripdock/

## SUPPLEMENTARY DATA

Experimental methods, Figures S1-S35 and Tables S1-S6 (PDF)

Supplementary Dataset 1: RiPPER analysis of lasso precursor peptides (XLSX)

Supplementary Dataset 2: RiPPER analysis of microviridin precursor peptides (XLSX)

Supplementary Dataset 3: RiPPER analysis of thiopeptide precursor peptides (XLSX)

Supplementary Dataset 4: RiPPER analysis of YcaO-TfuA precursor peptides (XLSX)

Supplementary Dataset 5: Phylogenetic tree of TfuA proteins and their association with peptides networks (PDF)

## ACKNOWLEDGEMENTS

We thank Bradley Moore (Scripps Institution of Oceanography, University of California San Diego, U.S.A.) for pCAP03, Vladimir Larionov (National Cancer Institute, NIH, U.S.A.) for *S. cerevisiae* VL6-48N, Mervyn Bibb (John Innes Centre, U.K.) for *S. coelicolor* strains, and Daniel Haft (NCBI/NIH, U.S.A.) for providing the precursor peptide HMMs. We thank Lionel Hill, Paul Brett and Gerhard Saalbach (John Innes Centre, Norwich, UK) for assistance with LC-MS, and Gwenaelle Le Gall and Ian Colquhoun (Quadrum Institute, Norwich, UK) for assistance with NMR.

