## Supplementary material for "Uncovering the unexplored diversity of thioamidated ribosomal peptides in Actinobacteria using the RiPPER genome mining tool"

### SUPPLEMENTARY METHODS

#### Chemicals

Unless otherwise specified, chemicals were purchased from Sigma-Aldrich, and enzymes from New England Biolabs. Molecular biology kits were purchased from Promega and GE Healthcare.

#### Strains and culture conditions

*Streptomyces varsoviensis* DSM 40346 was acquired from the German Collection of Microorganisms and Cell Cultures (DSMZ, Germany) and used as genetic source for the thiovarsolin gene cluster. *Streptomyces coelicolor* M1146, *S. coelicolor* M1152[1] and *Streptomyces lividans* TK21 were used as heterologous expression hosts. Unless otherwise specified, all *Streptom*yces strain were grown in SFM (solid) and TSB (liquid) media at 28 °C. Spores and mycelium stocks were kept at -20 °C and -80 °C in 20% glycerol. *Saccharomyces cerevisiae* VL6-48N[2] was used for transformation-associated recombination (TAR) cloning and was grown at 30 °C with shaking at 250 rpm in YPDA medium. Recombinant yeast selection was performed using selective media SD+CSM-Trp complemented with 5-fluoorotic acid (Fluorochem, 1 mg $mL^{-1}$). Yeast cell stocks were kept at -80 °C in 20% glycerol. *Escherichia coli* DH5α was used for standard DNA manipulations. *E. coli* DH5α BT340 was used for Flp-*FRT* recombination. *E. coli* BW25113/pIJ790 was used for Lambda-Red mediated recombination. *E. coli* ET12567/pR9604 and *E. coli* ET12567/pUZ8002 were used to transfer DNA to *Streptomyces* by intergeneric conjugation. All *E. coli* strains were grown in LB medium at 37 °C unless specified by particular protocols (pIJ790-carrying strains were grown at 30 °C for plasmid replication, and Flp-*FRT* recombination was performed at 42 °C). *E. coli* hygromycin selection was performed in DNAm (solid) and DNB (liquid) media. *E. coli* cell stocks were kept at -20 °C and -80 °C in 20% glycerol.

**Culture media composition**

All amounts given in g $L^{-1}$, unless otherwise specified:

**LB** (Lysogeny Broth): 10 tryptone, 5 yeast extract, 10 NaCl, pH 7. **DNB** / **DNAm** (Difco™ Nutrient Broth/Agar medium): purchased from Becton, Dickinson and Company. 20 agar was added for the solid version. **SFM** (Soya Flour Mannitol): 20 soya flour, 20 mannitol, 20 agar. **TSB** (Tryptic Soy Broth): Purchased from Oxoid. **BPM** (Bottromycin Production Medium): 10 glucose, 15 Difco™ soluble starch, 5 yeast extract, 10 soya flour, 5 NaCl, 3 $CaCO_3$. **MI** ("Monkey Island", modified actinonin production medium): 10 glucose, 10 Difco™ soluble starch, 20 corn liquor step, 20 soy flour, 2.5 $NH_4Cl$, 3 NaCl and 6 $CaCO_3$, pH 6.2. **TM1** (Teicoplanin Medium 1): 30 malt extract, 10 glucose, 15 soybean meal, 5 yeast extract, 4 $CaCO_3$. **E25**: 25 dextrose, 4 meat extract, 1 yeast autolysate, 10 soybean meal, 4 peptone, 2.5 NaCl, 5 $CaCO_3$. **GYM**: 4 glucose, 4 yeast extract, 10 malt extract, 2 $CaCO_3$, pH 7.2. **YPDA**: 10 bacto yeast extract, 20 bacto peptone, 20 glucose monohydrate, 40 mg $L^{-1}$ adenine hemisulfate. 20 agar were added for the solid version. **SD+CSM-Trp**: 1.7 YNB-AA-$(NH_4)_2SO_4$ (Formedium), 5 $(NH_4)_2SO_4$, 20 glucose, 20 agar, 20 adenine, and 740 mg $L^{-1}$ CSM-Trp (Formedium).

**TfuA-like protein retrieval and phylogenetic analysis**

The NCBI Conserved Domain Architecture Retrieval Tool[3] (CDART) was used to retrieve all TfuA domain protein sequences from the phylum Actinobacteria in the NCBI non-redundant protein sequence database. These 325 proteins were manually assessed by Pfam analysis for TfuA domains, which resulted in the removal of five proteins from this dataset. To limit the overrepresentation of highly similar proteins in an analysis of phylogeny and gene cluster diversity, ElimDupes (https://www.hiv.lanl.gov/content/sequence/ELIMDUPES/elimdupes.html) was used to remove proteins with at least 99% identity to each other from the dataset to leave one representative protein. This provided a dataset of 229 TfuA domain proteins. Three proteins that contained fused YcaO and TfuA domains were removed for phylogenetic analysis, along with one (KZS83678.1) that is truncated. The standalone TfuA domain protein dataset (225 proteins) was aligned using MUSCLE 3.8.31[4] with default settings. The resulting alignment was used to construct a maximum likelihood tree using RAxML-HPC2 on XSEDE (with 100 bootstrap replications) on the CIPRES Science Gateway (https://www.phylo.org/). The tree was visualized using the interactive Tree Of Life[5] (iTOL) (Supplementary Dataset 5).

**RiPPER details**

RiPPER consists of a series of Perl script that require the RODEO2 Python script,[6,7] BioPerl,[8] a locally installed Pfam database[9,10] and a modified build of Prodigal[11] (Prodigal-short) to operate. Analysis parameters for RiPPER are defined in an associated configuration file (local.conf), which can be modified to optimize the genome mining process. EGN (Evolutionary Gene and genome Network)[12] was used to construct protein similarity networks, which were visualized using Cytoscape 2.8.3.[13] Further information is provided in the documentation provided with the RiPPER scripts at https://github.com/streptomyces/ripper. For ease of use, a Docker container is provided that contains all features required for using RiPPER. This is available at https://hub.docker.com/r/streptomyces/ripdock/ along with instructions on installation and usage. A workflow for using RiPPER is described below.

**Workflow for RiPPER**

Below is a summary of the RiPPER workflow, which has been developed for gene cluster visualization in Artemis[14] (Figure S1). Where relevant, default analysis parameters are listed. These are all customizable from the local.conf configuration file associated with a given RiPPER analysis; variables are highlighted below in bold.

1. Using RODEO,[6,7] accession numbers for a set of putative RiPP tailoring enzymes (RTEs) are used to obtain nucleotide regions (as GenBank files) centered on the tailoring enzyme, which is highlighted as a green gene for clarity in Artemis. 25 kb regions were obtained for the TfuA analysis (**flankLen** = 12.5 kb), and 35 kb regions were obtained for the known RiPP families (**flankLen** = 17.5 kb, default).
2. Every retrieved genomic region is subjected to RODEO analysis to obtain a RODEO output for each input accession, as well as Pfam domain data across the gene cluster.
3. GenBank files are then analyzed using a specially built version of Prodigal,[11] which we call Prodigal-short. This is configured to find genes as short as 60 nucleotides instead of the usual size cut-off of 90 nucleotides.
4. For all the genes found by Prodigal-short the following is done:
    - **a.** The Prodigal score is enhanced if the gene is on the same strand as the tailoring enzyme (**sameStrandReward**, default = 5).
    - **b.** Genes are only retained for analysis if they overlap with existing annotated genes by 20 nucleotides or less.
    - **c.** RiPPER uses Prodigal-short to only identify putative ORFs within a likely size window for precursor peptide genes. Therefore, genes are only retained for analysis if the length of the encoded peptide is between **minPPlen** and

**maxPPlen**. A window of 20 – 120 AA (default) was used in all analyses in this study.

  d. If a gene is not filtered out in the above steps, it is annotated in the GenBank file and its distance from the tailoring enzyme is determined.
  e. All putative genes identified are provided in the resulting GenBank file and are color-coded from pale red (low score) to bright pink (high score) (Figure S1). Scoring criteria are viewable in Artemis as notes for each putative gene.
  f. RiPPER also retrieves and scores genes that were already annotated if they encode peptides below the **maxPPlen** (default = 120 AA). This means that annotated precursor peptides are also retrieved for downstream analysis.

5. The resulting annotated GenBank files can be viewed in Artemis at this stage for manual identification of RiPP precursor peptides.
6. If the gene is within a specified distance (**maxDistFromTE**) from the RTE, it is included in the output list and also saved in a Sqlite3 table. A distance of ± 8 kb is used as default.
7. Within this region, the top scoring short peptides (no lower score threshold) are retrieved. The number retrieved is defined by **fastaOutputLimit** (default = 3) In addition, any further peptides with Prodigal scores over a threshold (**prodigalScoreThresh**) within this region are retrieved. A score threshold of 15 was used in the TfuA analysis and a score threshold of 7.5 (default) was used in the analysis of known RiPP families.
8. All retrieved peptides are analyzed for Pfam domains, and all information is tabulated alongside various associated data (tailoring enzyme accession, strain, peptide sequence, distance from tailoring enzyme, coding strand in relation to tailoring gene, Prodigal score) in a tab-separated out.txt file. All data are collated in a single file if multiple genomic regions are analyzed in parallel.
9. All peptides identified by RiPPER across the entire Genbank file that were not retrieved in step 8 (no distance or score threshold) are searched for characterized precursor peptide domains.[10] Data for these peptides is then tabulated in a tab-separated distant.txt file.
10. **Optional follow-on analysis: protein similarity networking and BGC comparative analysis.** Protein similarity networking does not form part of the automated RiPPER workflow, but this does assist with the identification of authentic precursor peptides. The RiPPER output includes fasta files (out.faa and distant.faa) for all retrieved peptides that are compatible for analysis with EGN.[12] The following settings were used for all analyses: E-value threshold = 10, hit identity threshold = 40%, hit covers at least 35% of the shortest sequence, minimum hit length = 15 AA. The resulting networks

were visualized using Cytoscape 2.8.3,[13] where data obtained from RiPPER were imported as node attributes. The similarity between BGCs associated with the same network was assessed using MultiGeneBlast.[15] Peptides from each network were aligned using MUSCLE[4] and alignments were visualized using ESPript 3.0.[16]

**Statistical analysis of predicted precursor peptides**

The RiPPER output indicates whether a predicted peptide-coding ORF was already annotated in the original GenBank file. Data for predicted precursors peptides in BGCs identified from the top 30 networks (Networks 1, 2, 3, 5, 7, 9, 12, 15, 20, 21, 22, 24, 26, 27 and 28; Table S1) were retrieved and classified as "Not annotated" (88 peptides) or "Already annotated" (73 peptides) (Figure S20). The statistical significance of the difference in peptide length distributions between these sample sets was assessed using the two-tailed Mann-Whitney *U* test, which provided a *p*-value of <0.0001 (Z-score = -9.04956).

**Identification of precursors to lasso peptides, microviridins and thiopeptides**

Studies by Tietz *et al.*,[6] Ahmed *et al.*,[17] and Schwalen *et al.*[7] had previously used RiPP tailoring enzyme accessions to mine for precursors to lasso peptides, microviridins and thiopeptides, respectively. The same accession codes were used to mine for precursor peptides using RiPPER (Supplementary Datasets 1-3), although not all accessions could be retrieved as some records no longer exist on NCBI. RiPPER was run using analysis parameters as described above and the results are described in Table 1. Peptide similarity networking was carried out using EGN (as described above), which provided large networks for each dataset (Network 1, Figures S2-S4, Supplementary Datasets 1-3). To determine the ability of RiPPER to retrieve authentic precursor peptide sequences, a bespoke script was used to compare the RiPPER outputs with the prior studies.

**Transformation-associated recombination (TAR) cloning and heterologous expression of the thiovarsolin gene cluster**

A vector to capture the thiovarsolin gene cluster from *S. varsoviensis* genomic DNA (gDNA) was constructed using yeast assembly between a linearized pCAP03 vector[18] and two single-strand oligonucleotides (TARvar-1 and TARvar-2). Oligonucleotides had 35 nucleotide homology sequences with pCAP03 and were designed to generate a vector with 50 nucleotide homology sequences with upstream and downstream regions of the gene cluster either side of a PmeI restriction site. pCAP03 was digested with XhoI and NdeI, and the linearized plasmid and ss-oligos (1:10 ratio) were transformed into *S. cerevisiae* VL6-48N by lithium acetate/polyethylene glycol 3350 mediated transformation. For yeast-colony PCR, each colony was resuspended in 50 µL 1 M sorbitol (Fisher) and 2 µL of zymolyase (5 U $\mu L^{-1}$) added to each cell suspension and incubated at 30 °C for 1 hour. Cell suspensions were then boiled for 10 minutes, centrifuged (15 s, 1,000 x *g*) and 1 µL of the supernatant was analyzed by PCR.

To transfer the plasmids from yeast into *E. coli*, colonies of yeast were grown in 10 mL of liquid SD+CSM-Trp for 18 h at 250 rpm, 30 °C. Cells were harvested by centrifugation (5 min, 1,789 x *g*), and resuspended in 200 µL 1 M sorbitol plus 2 µL of zymolyase (5 U $µL^{-1}$). Cell suspensions were incubated at 30 °C for 1 hour to produce spheroplasts, which were then pelleted (10 min, 600 x *g*). The supernatant was aspirated, and plasmid DNA extracted from the pellet using a standard Wizard miniprep protocol (Promega). 1 µL plasmid DNA was then transformed into *E. coli* DH5α by electroporation and selected with kanamycin (50 µg $mL^{-1}$) Colonies containing the correct capture vector were identified by PCR (primers: CAP03_check-fw and CAP03_check-rv), and the plasmid was isolated and confirmed by sequencing.

gDNA from *S. varsoviensis* was digested with EcoRV and ScaI, and the pCAP03-derived capture vector was linearized between the capture arms with PmeI. These were both then introduced into *S. cerevisiae* VL6-48N by spheroplast polyethylene glycol 8000 transformation. Successful gene cluster capture by pCAP03 was confirmed by colony PCR (primers: TARcheck-fw and TARcheck-rv). The plasmids from three positive clones were recovered and transformed into electrocompetent *E. coli* DH5α for amplification and further restriction analysis of the purified construct (pTARvar). *E. coli* ET12567/pR9604 was transformed with pTARvar by electroporation, and transformants were then used to transfer pTARvar into *S. coelicolor* (M1146 and M1152) and *S. lividans* TK21 by intergeneric conjugation. Nalidixic acid (25 µg $mL^{-1}$) and kanamycin-resistant (50 µg $mL^{-1}$) exconjugants containing integrated pTARvar (*S. coelicolor* M1146-var, *S. coelicolor* M1152-var and *S. lividans* TK21-var) were verified by PCR using GoTaq polymerase (Promega) (primers: TAR_check-fw and TAR_check-rv).

**Fermentation conditions for metabolite screening**

Seed cultures of *S. coelicolor* M1146-var, *S. coelicolor* M1152-var and *S. lividans* TK21-var were obtained by fermentation in a 250 mL flask containing 50 mL of TSB for 72 h. 250 µL seed culture was used to inoculate 5 mL of a variety of culture media (TSB, BPM, GYM, MI, TPM, E25; see Culture media composition section) in 50 mL conical centrifuge tubes with caps replaced by foam bungs. Control strains carrying a genome-integrated empty pCAP03 vector were cultured in the same way for comparison. All fermentations were conducted in triplicate and incubated at 28 °C with shaking at 250 rpm. Culture samples (500 µL) were taken at 72 h and 168 h, mixed with one volume of methanol and agitated for 30 min at room temperature. These mixtures were then centrifuged (13,000 rpm, 30 min) and 600 µL of the resulting supernatant was transferred to glass vials for liquid chromatography-mass spectrometry (LC-MS) analysis.

**LC-MS analysis**

Spectra were obtained using a Shimadzu Nexera X2 UHPLC coupled to a Shimadzu IT-TOF mass spectrometer. Samples (5 µL) were injected onto a Phenomenex Kinetex 2.6 µm XB-C18 column (50 mm x 2.1 mm, 100 Å) set at a temperature of 40 °C and eluting with a linear gradient of 5 to 95% acetonitrile in water + 0.1% formic acid over 6 minutes with a flow-rate of 0.6 mL $min^{-1}$. Positive mode mass spectrometry data was collected between *m/z* 200 and 2000, and $MS^2$ data was collected using collision-induced dissociation of the most abundant singly charged species in a scan, with an exclusion time of 0.8 seconds. Untargeted comparative metabolomics was carried out on triplicate data using Profiling Solution 1.1 (Shimadzu) with an ion *m/z* tolerance of 100 mDa, a retention time (RT) tolerance of 0.1 min and an ion intensity tolerance of 100,000 units.

For the accurate mass measurement of the thiovarsolins, high-resolution mass spectra were acquired by LCMS on a Synapt G2-Si mass spectrometer equipped with an Acquity UPLC (Waters). Samples were injected onto an Acquity UPLC® BEH C18 column, 1.7 µm, 1x100 mm (Waters) and eluted with a gradient of (B) acetonitrile/0.1% formic acid in (A) water/0.1% formic acid with a flow rate of 0.08 mL $min^{-1}$ at 45 °C. The concentration of B was kept at 1% for 2 min followed by a gradient up to 30% B in 4 min. MS data were collected with the following parameters: resolution mode, positive ion mode, scan time 0.5 s, mass range *m/z* 50-1200 (calibrated with sodium formate), capillary voltage = 3.0 kV; cone voltage = 40 V; source temperature = 120 °C; desolvation temperature = 350 °C. Leu-enkephalin peptide was used to generate a lock-mass calibration with *m/z* = 556.2766 measured every 30 s during the run.

**Deletion of genes in the thiovarsolin biosynthetic gene cluster**

The mutational analysis of the thiovarsolin BGC was performed using an *E. coli*-based Lambda-Red-mediated PCR-targeting strategy,[19] which allowed the substitution of genes or groups of genes in pTARvar by a PCR-generated cassette containing the apramycin resistance gene *aac(3)IV*. Given the presence of an *oriT* in the original pCAP03 vector, the upstream primer design was modified with respect to the original protocol in order to exclude a second *oriT* from the PCR-targeting resistance cassette and avoid undesired recombinations. Therefore, resistance cassettes were PCR amplified using pIJ773 as template (see primers in Table S3 and mutants in Tables S3 and S5). In the case or *varA*, and additional in-frame deletion mutant affecting only the core precursor peptide was created employing a pIJ773-derived cassette lacking OriT (pIJ773 Δ*oriT*) but preserving both *FRT* recombination sites (primers RD1 and RD3), which allowed the elimination of the apramycin resistance cassette after Flp-*FRT* recombination in *E. coli* DH5α BT340 and the creation of a clean *varA* mutant (Δ*varA*_clean). The PCR-targeting mutant versions of pTARvar were transferred to *S.*

*coelicolor* M1146 by *E. coli* ET12567/pUZ8002-mediated intergeneric conjugation and selected by resistance to nalidixic acid (25 µg $mL^{-1}$), kanamycin (50 µg $mL^{-1}$) and, when required, apramycin (50 µg $mL^{-1}$).

Constructs for the complementation of mutants showing differences in thiovarsolin production in comparison to *S. coelicolor* M1146-var (Δ*varA*, Δ*varY*, Δ*varT* and Δ*varO*) were obtained by high-fidelity PCR amplification (Herculase II, Agilent) of each of these genes (primers CP1 and CP2 for *varA*, CP3 and CP4 for *varA*p, CP5 and CP6 for *varY*, CP7 and CP8 for *varT*, and CP9 and CP10 for *varO*), digestion of the PCR product with NdeI and HindIII and cloning by ligation (T4 DNA ligase, Invitrogen) into NdeI – HindIII digested pIJ10257.[20] Ligation mixtures were transformed into chemically competent *E. coli* DH5α, plasmids were recovered by alkaline lysis and then sequenced. The resulting plasmids (pJ10257-*varA*, pIJ1027-*varA*p, pIJ10257-*varY*, pIJ10257-*varT* and pIJ10257-*varO*) were introduced into the corresponding *S. coelicolor* M1146-var mutants by *E. coli* ET12567/pUZ8002-mediated intergeneric conjugation. Exconjugants were selected by resistance to nalidixic acid (25 µg $mL^{-1}$), kanamycin (50 µg $mL^{-1}$), hygromycin (50 µg $mL^{-1}$) and, when required, apramycin (50 µg $mL^{-1}$).

**Construction of a minimal thiovarsolin gene cluster (pIJ10257-*varA*p*YT*)**

High-fidelity PCR amplification (primers CP3 and CP8) was used to obtain a 3.7 kb DNA fragment containing *varA* (including its putative promoter region), *varY* and *varT*. The gel-purified PCR product was digested with NdeI and HindIII, and cloned by ligation into NdeI - HindIII digested pIJ10257. The resulting construct (pIJ10257-varAYT) was introduced into *S. coelicolor* M1146 by intergeneric conjugation and selection with nalidixic acid (25 µg $mL^{-1}$) and hygromycin (50 µg $mL^{-1}$).

**Site-directed mutagenesis of *varA***

The *varA* gene was mutated to generate a version that encodes a peptide where the four repetitions containing APR were replaced by repetitions containing GPR, and residues flanking this motif were mutated to be identical to the natural GPR-containing repeat (peptide VarA*, Figure S35). The presence of highly repetitive sequences and high G+C content meant that commercial gene synthesis failed. Therefore, the construction of the *varA** mutant was achieved by a series of primer extension PCR reactions employing codon degenerated primers to avoid undesired annealing in the repetitive core region of *varA*. To achieve multiple mutations across such a highly repetitive peptide, this method necessitated the introduction of two additional mutations within the third repeat region (Figure S35).

An initial DNA fragment (including the *varA* promoter region, the leader peptide and the first 2.5 mutant repetitions of the core peptide) was generated by three consecutive PCR extension reactions (first primers: CP3 and AG1; second primers: CP3 and AG2; third primers: CP3 and AG3), which was gel purified, digested with NdeI and XbaI, and ligated into NdeI - XbaI linearized pGP9[21] to yield pGP9_varA*p_partial. A second DNA fragment (containing the last 2.5 mutated repetitions of the core peptide and the intergenic region following *varA*) was generated by three consecutive PCR extension reactions (first primers: AG4 and CP4; second primers: AG5 and CP4; third primers: AG6 and CP4). This was gel purified, digested with KpnI and Hind III, and cloned into KpnI – HindIII linearized pGP9_*varA**_partial to yield the final construct, pGP9_*varA**p, which was sequenced and introduced into *S. coelicolor* M1146-var Δ*varA*_clean by intergeneric conjugation and selection with nalidixic acid (25 μg $mL^{-1}$), kanamycin (50 μg $mL^{-1}$) and apramycin (50 μg $mL^{-1}$).

**Large scale fermentation, isolation and structural elucidation of thiovarsolins A and B**

6 x 2 L flasks, each containing 500 mL BPM, were inoculated with 25 mL of *S. coelicolor* M1146_var seed culture grown in TSA (200 mL in a 1 L flask, 72 h) and incubated at 28 °C with shaking at 250 rpm for 10 days. The culture broth was separated from the mycelium by centrifugation to yield a cell-free supernatant (ca. 3 L), which was concentrated to dryness to afford 32.4 g of crude extract. This extract was resuspended in 0.4 L of distilled water and was then fractionated using vacuum liquid chromatography (VLC) on C-18 RP silica gel using a gradient of $H_2O$:MeOH (100:0 to 0:100). The major thiovarsolin-containing fraction (2.4 g) was subsequently fractionated on a Sephadex LH-20 column using methanol:water (7:3) as the mobile phase. Thiovarsolin-containing fractions were combined and dried to yield 0.78 g. This was subjected to solid phase chromatography using a C-18 cartridge (Discovery DSC-18, 20 mL) with a gradient of water:methanol (100:0 to 70:30). Fractions containing thiovarsolins A and B were further purified by semipreparative HPLC (Phenomenex Luna PFP(2), 210 mm x 10 mm, 5 mm; 3 mL $min^{-1}$, UV detection at 268 nm) with a linear gradient of $CH_3CN/H_2O$ (+0.1% formic acid) from 5 to 15% $CH_3CN$ over 30 minutes yielding thiovarsolin A (1.3 mg, retention time = 15.2 min) and thiovarsolin B (0.8 mg, retention time = 18.3 min). 1D and 2D NMR spectra were recorded on Bruker Avance 700, 600 and 400 MHz NMR spectrometers operated using Topspin 2.0 software. Spectra were calibrated to the residual solvent signals of DMSO-$d_6$ with resonances at $\delta_H$ 2.50 and $\delta_C$ 39.52.

### SUPPLEMENTARY FIGURES

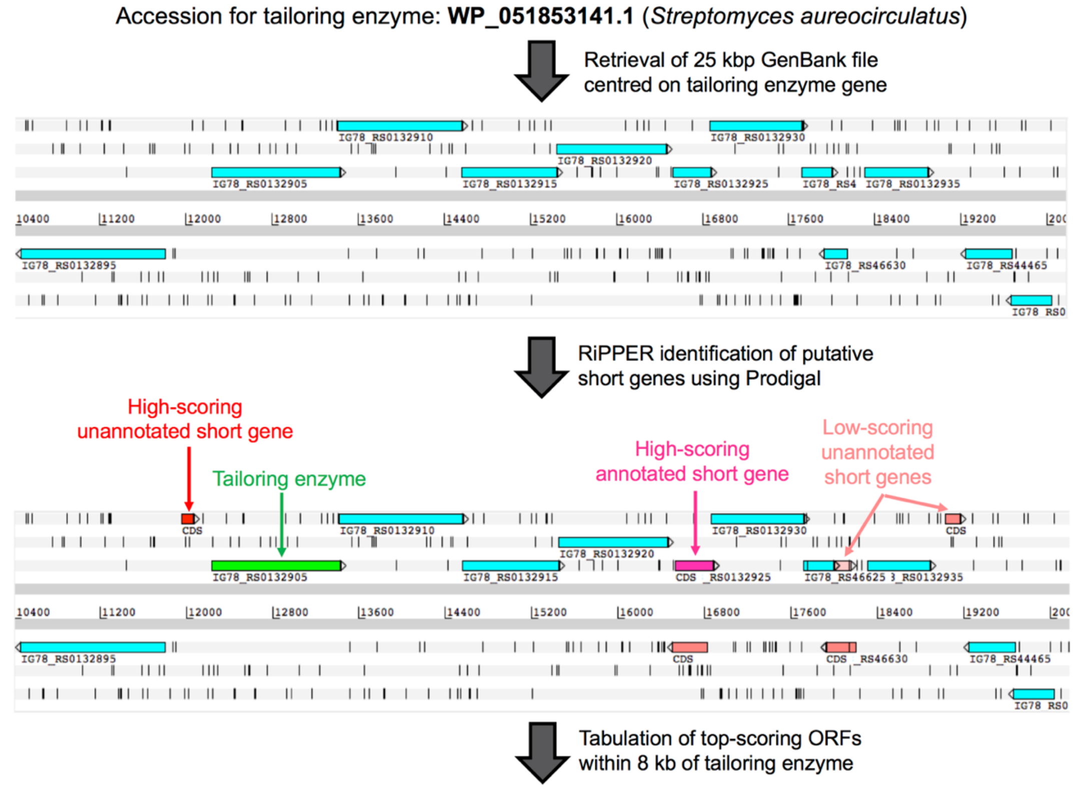


| Accession | Organism | PP Serial | FastaID | Sequence | Same Strand | PP_TE Distance | Prodigal score | Pfam |
|---|---|---|---|---|---|---|---|---|
| WP_051853141 | *Streptomyces aureocirculatus* | 1 | WP_051853141_1 | MHASVLSAVEIALTLALVAWMATGFTRLPGRWLRWKMFCRATFTIV TLTGTKDGRTEPVNVYDHLSPGSFILGPPQLQTILNYLTHTGRYDR VDGHGRLLNADGERPIDIRDSRVVL | yes | 3100 | 62.07 | none |
| WP_051853141 | *Streptomyces aureocirculatus* | 2 | WP_051853141_2 | MVSKSQAAVSTLKPTQETEELTDTVVVAVDVEDEE | yes | 174 | 18.35 | none |
| WP_051853141 | *Streptomyces aureocirculatus* | 3 | WP_051853141_3 | MLEPSGSNNRGQVPTNVSRLNGRVPARPAKDEPEHYHLDIGYAFT TAHADVGSIQEAEVTGAAWHQLAEAESLVGSRIARAVSTPARTD | no | 4501 | 1.95 | none |

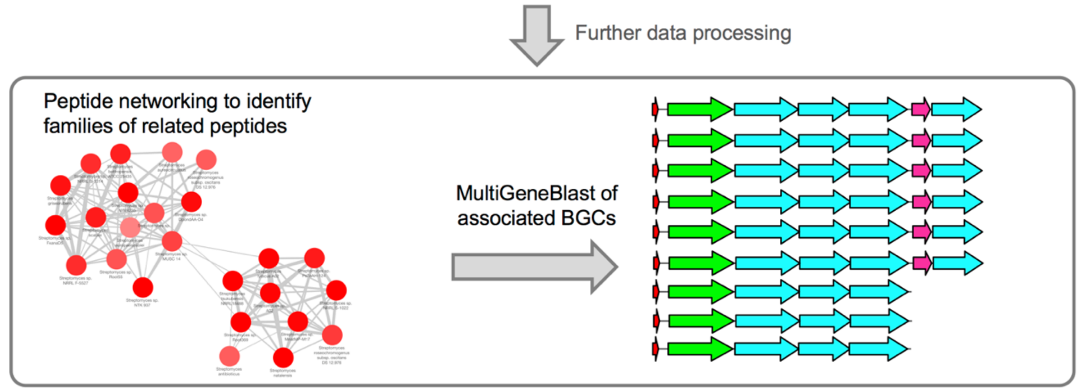


**Figure S1.** RiPPER workflow showing the output of a single TfuA accession (a short region of the Artemis[14] output is shown), as well as the associated peptide network and MultiGeneBlast[15] analysis (the output is simplified for clarity and simply reflects the Artemis color-coding).

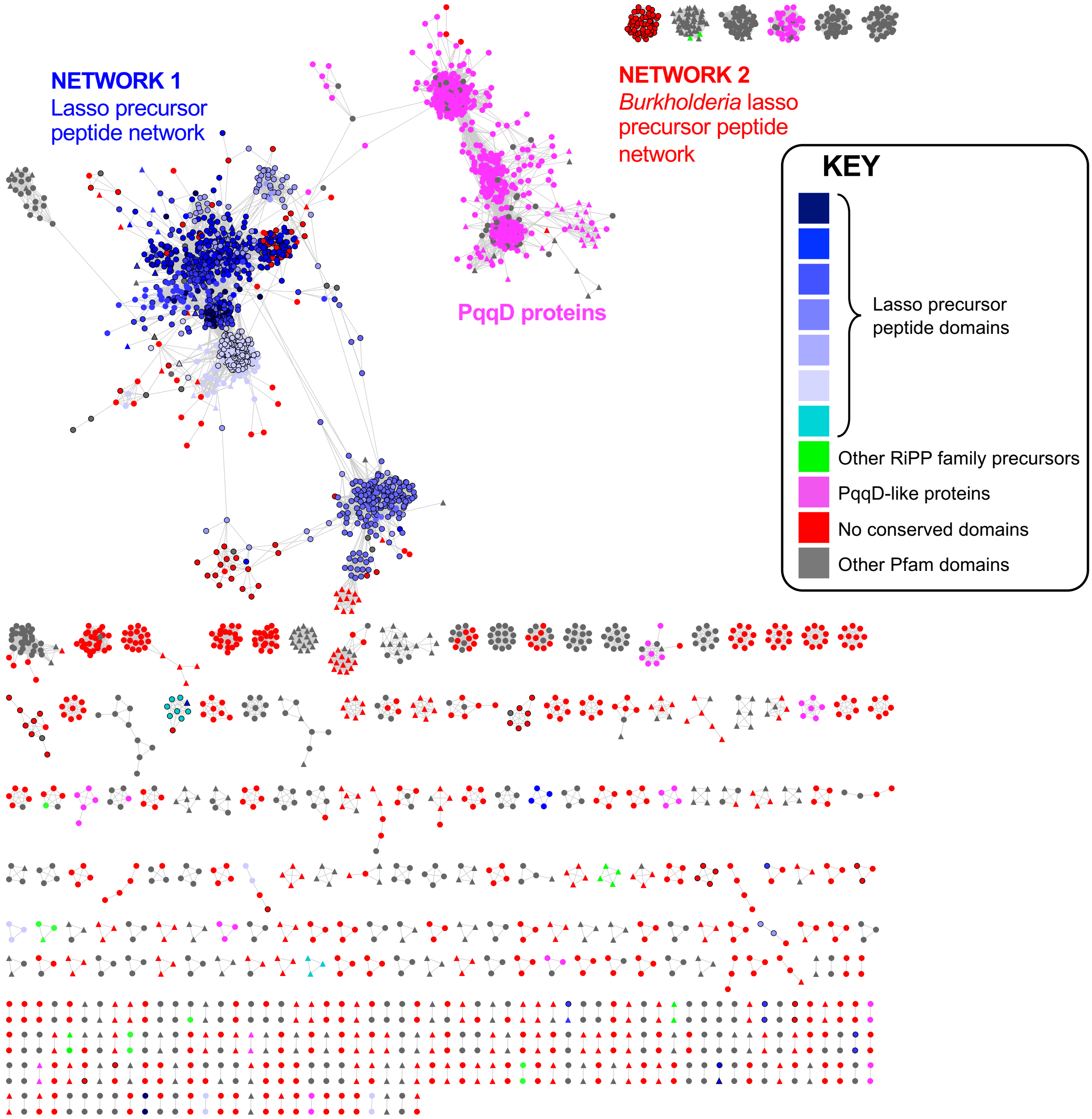


**Figure S2.** Peptide similarity networks of short peptides identified by RiPPER that are encoded in BGCs of asparagine synthetases previously predicted by RODEO to catalyze lasso peptide formation.[6] Each node represents one peptide, have a black border if they match a peptide identified by Tietz *et al.*,[6] and are colored by Pfam domain. Putative peptides encoded on the same strand as the asparagine synthetase gene have circular nodes and those on the opposite strain have triangular nodes. An edge cut-off of 40% identity was used.

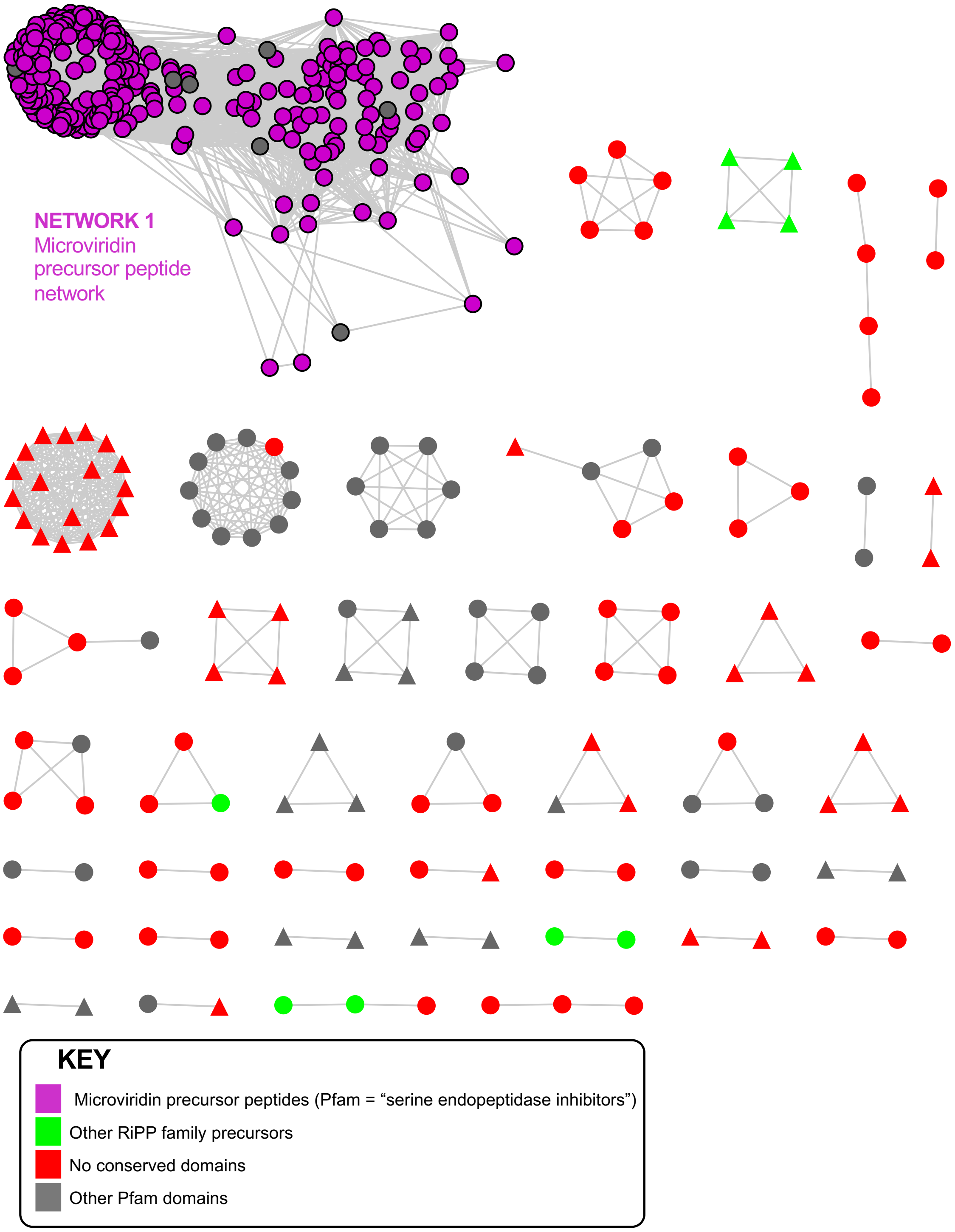


**Figure S3.** Peptide similarity networks of short peptides identified by RiPPER that are encoded in BGCs encoding homologues of MvdD (or MvdC) that were previously predicted by Ahmed *et al*. to produce microviridins. [17] Each node represents one peptide, have a black border if they match a peptide identified by Ahmed *et al*.,[17] and are colored by Pfam domain. Putative peptides encoded on the same strand as the *mvdD/C* gene have circular nodes and those on the opposite strain have triangular nodes. An edge cut-off of 40% identity was used.

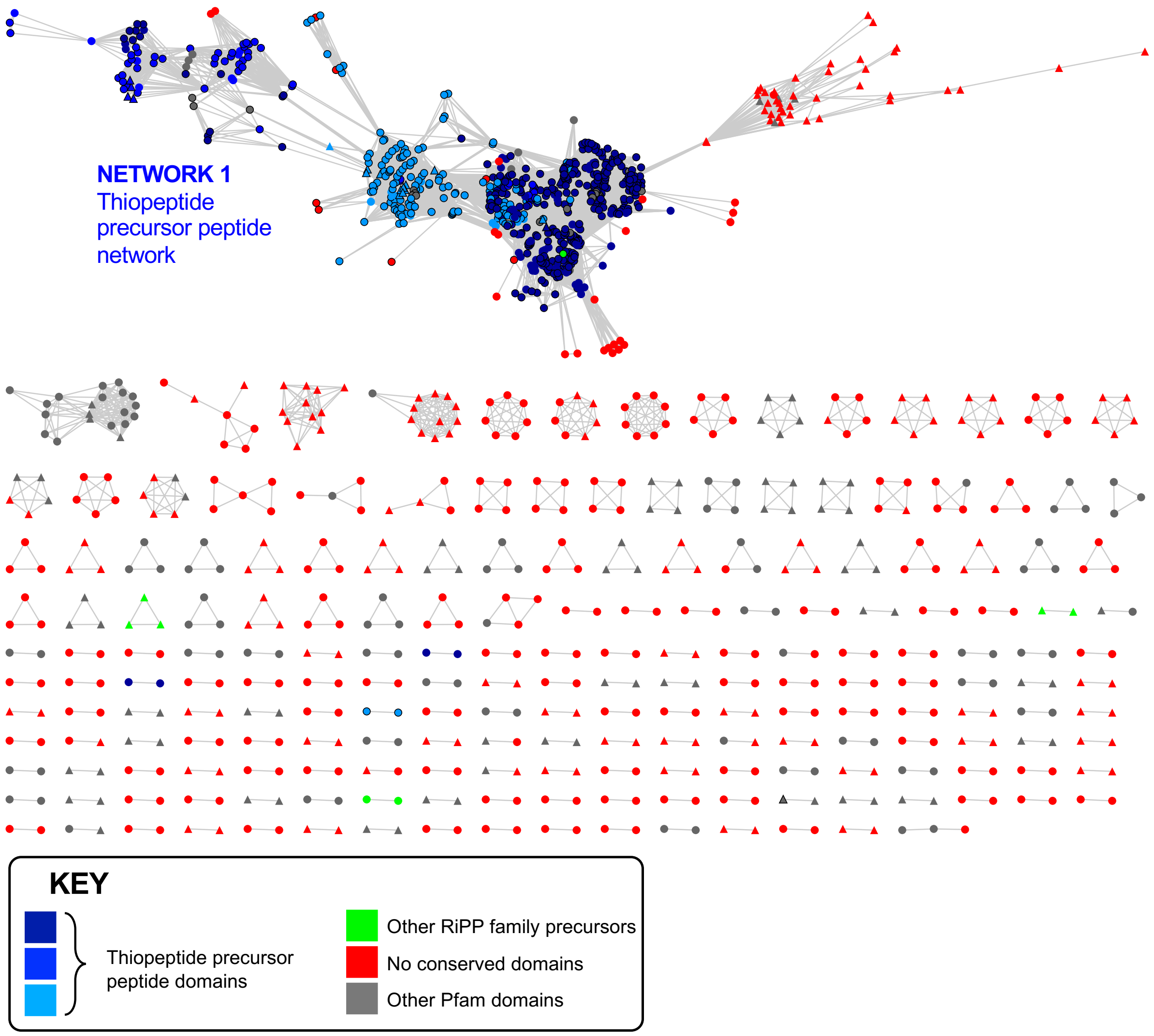


**Figure S4.** Peptide similarity networks of short peptides identified by RiPPER that are encoded in BGCs encoding [4 + 2]-cycloaddition enzymes that were previously predicted by Schwalen *et al*. to produce thiopeptides.[7] Each node represents one peptide, have a black border if they match a peptide identified by Schwalen *et al*.,[7] and are colored by Pfam domain. Putative peptides encoded on the same strand as genes encoding [4 + 2]-cycloaddition enzymes have circular nodes and those on the opposite strain have triangular nodes. An edge cut-off of 40% identity was used.

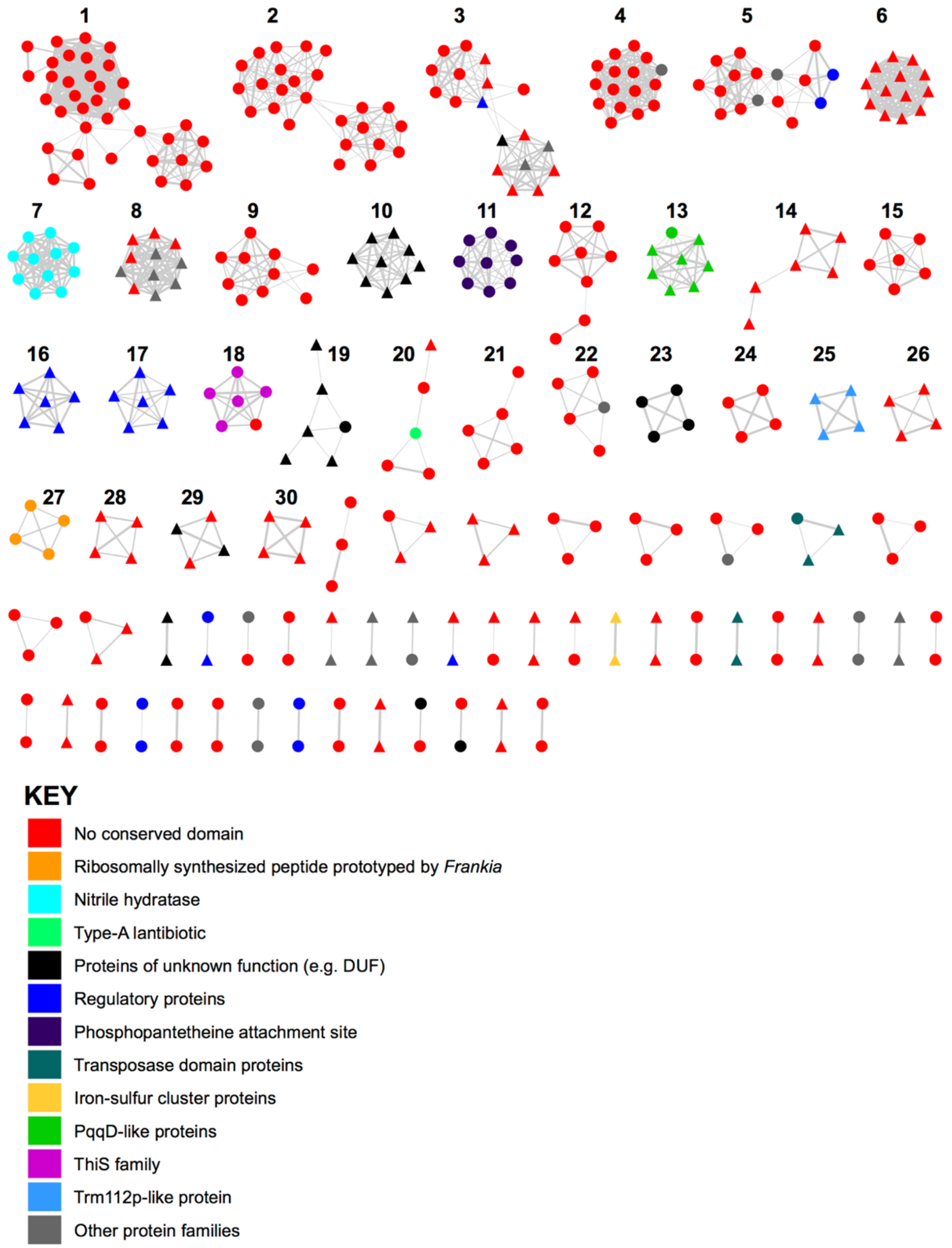


**Figure S5.** All networks identified for short peptides associated with *tfuA*-like genes using EGN[12] with an identity cut-off of 40%. The top 30 networks (4 or more peptides) are numbered and peptides are color-coded according to Pfam domain. Each node represents a single peptide and are circular if they are on the same strand as the *tfuA*-like gene and triangular if they are on the opposite strand.

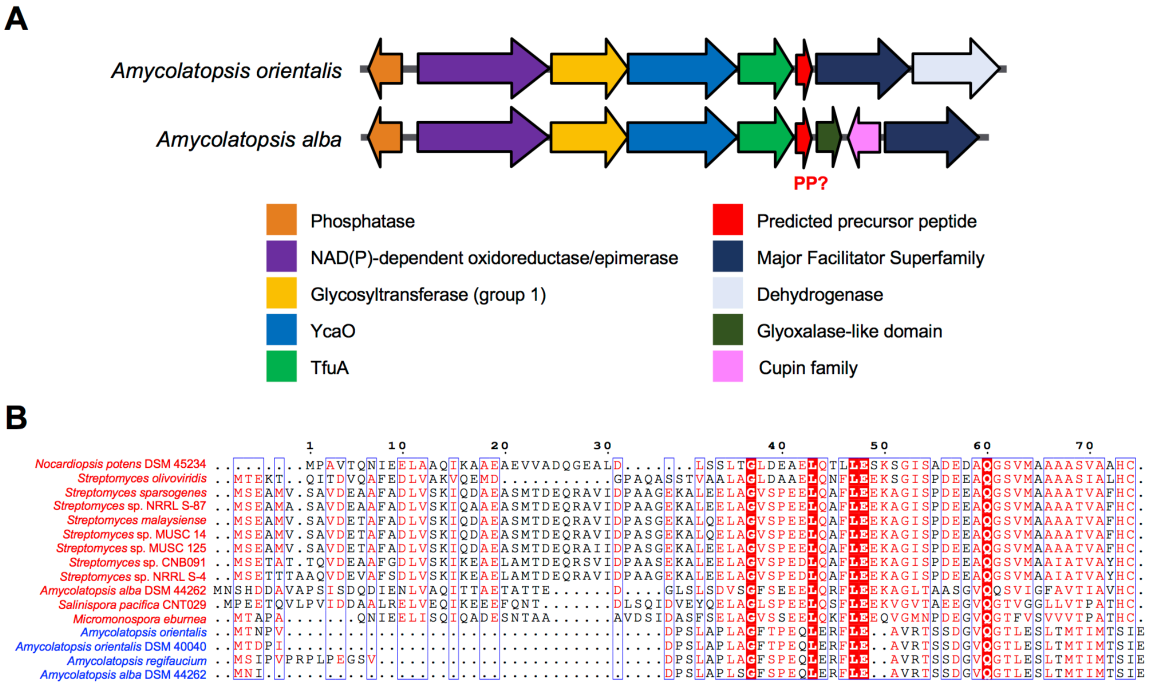


**Figure S6.** Details of the thioviridamide-containing Network 5. (A) Representative conserved BGCs relating to Fig. 3 in main paper. (B) Alignment of the thioviridamide-like precursor peptides that form Network 5. Peptides with red names indicate peptides with thioviridamide-like features (e.g. terminal HC motif) and BGCs with thioviridamide-like biosynthetic genes, whereas peptides with blue names reflect networked peptides with different putative core sequences and BGCs lacking almost all thioviridamide-like biosynthetic genes. See Frattaruolo *et al.*[22] for details of thioviridamide-lik BGCs. All alignment images were generated using Espript 3.0.[16]

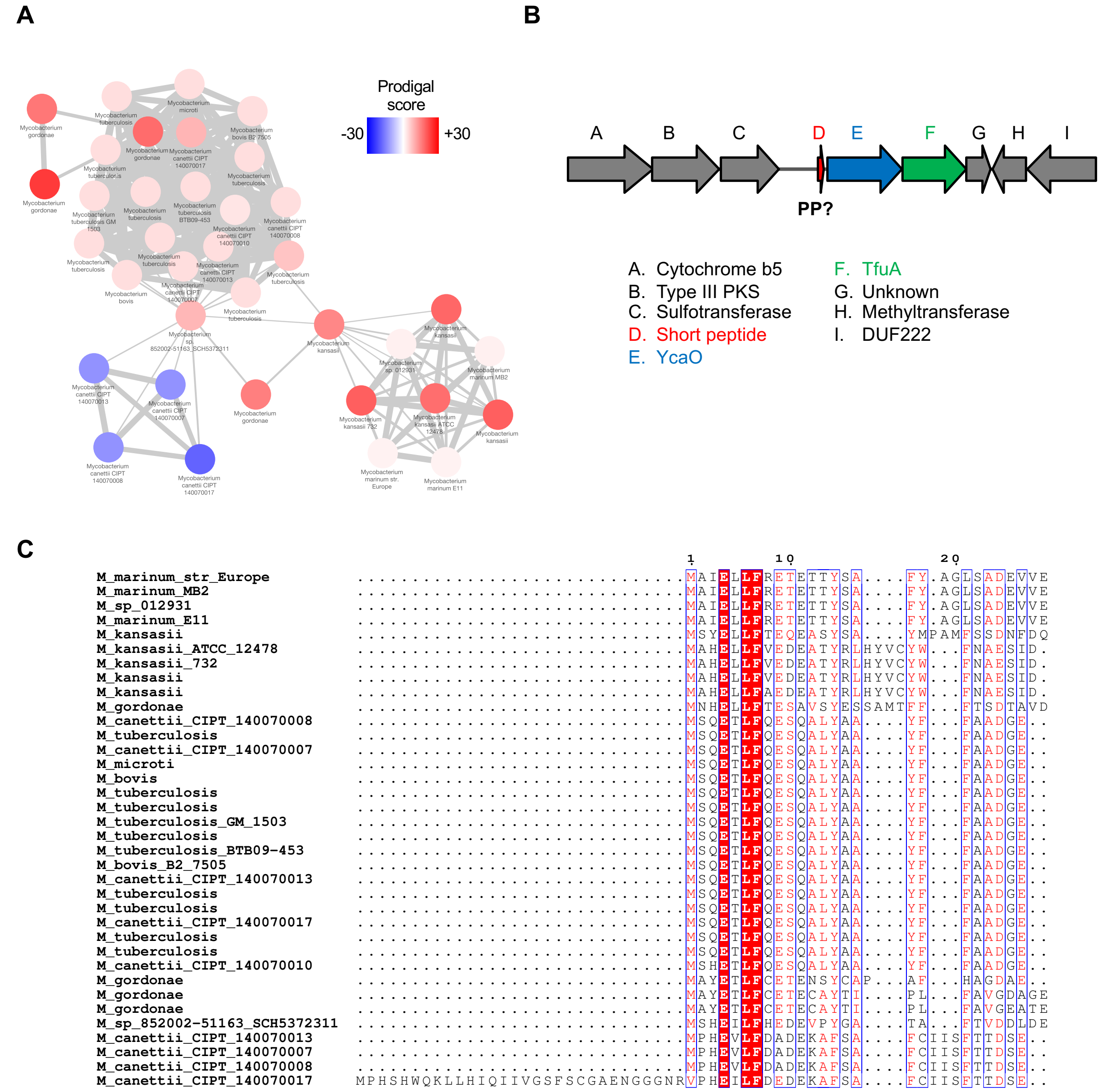


**Figure S7.** Network 1 overview. (A) Peptide network color-coded by Prodigal score (scores over 30 are all colored red). (B) Representative conserved BGC with PP, YcaO and TfuA genes highlighted. (C) Sequence alignment of all peptides present in network. A red background indicates full conservation, blue boxes represent positions with over 60% sequence equivalence, with the corresponding similar amino acids colored red.

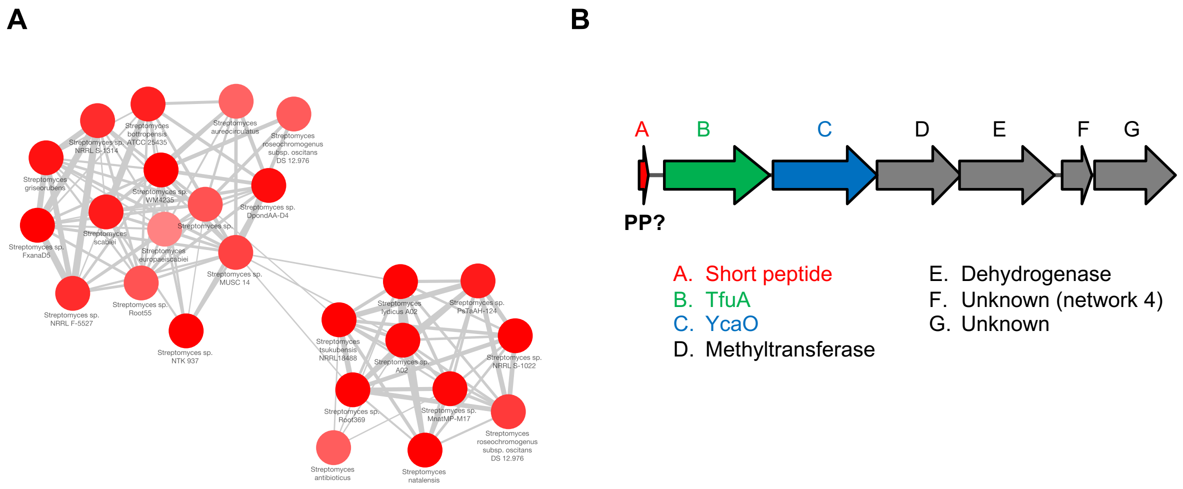


C

```
                                                         1         10          20         30         40
Streptomyces_antibioticus          ..........MEHLSKSETVA...AAQQTT...TVRLADIAVPSGTVRIEDVEDEGP...
Streptomyces_europaeiscabiei       MRATGPHRWNQHTREKSVESKSRAAAISNA...SVDVPEDITTELVVVAIDVDDEE....
Streptomyces_sp_NRRL_S-1022        ............MVSKAQDAA.QATEQVAD...GADVEALVDIAAVVRSEDVEQEECYTF
Streptomyces_roseochromogenus      .............MSKAQEAA.QSTAVVAD...GTDVEELVDTSAVVRSEDVEQDECYTF
Streptomyces_sp_PsTaAH-124         ............MISKTQEAA.QQTVPVAD...PTDVAALVDTSAVVRSEDVEQEECYTF
Streptomyces_sp_Root369            .............MNKAQEAA.NTTPAVAD...ADDVAALVDTSAVVRSEDVEQEECYTF
Streptomyces_tsukubensis           ...........MEQAKSQLVA...QAAEIQ...EIDVADLVDPSAIVQSEDVEQEECYTF
Streptomyces_sp_MnatMP-M17         ...........MDLSKSQVAAVQNSGQEGDTSVATDVTELIEPSAVVRSEDTEQDECYTF
Streptomyces_natalensis            ............MEAKSQVIA.QAVGPAIE...ETDVVDLIDPAKVVKSEDVEQEECFTF
Streptomyces_sp_A02                ...........MDIAKSQVAA.ESSGQQIE...ETNVADLIDPSAVVRSEDVEQEECFTF
Streptomyces_lydicus_A02           ...........MDIAKSQVAA.ESSGQQIE...ETNVADLIDPSAVVRSEDVEQEECFTF
Streptomyces_sp                    ...........MEQSKSAIAA...ANSPIP...AHEAADITRP..VVIAVDVEDEE....
Streptomyces_griseorubens          ............MQFESTSAA..QSAVQAP...AREPDEVTGP..VVVAVDVEDEE....
Streptomyces_sp_NRRL_S-1314        ...........MTQSKSTRAA..MSAVQAP...AREPDEVTGP..VVVAVDVEDEE....
Streptomyces_sp_NRRL_F-5527        ...........MTQSKSTRAA..MSAVQAP...AREPDEVTGP..VVVAVDVEDEE....
Streptomyces_sp_FxanaD5            ...........MTQSKSTSAA..MSAVQAP...AREPDEVTGP..VVVAVDVEDEE....
Streptomyces_scabiei               ............MEAKSQTAA......RTS...TTQTPENLATEEVVIAVDVDDEE....
Streptomyces_bottropensis          ............MEAKSQAAA......RTS...TTQTPENLATEEVVIAVDVDDEE....
Streptomyces_sp_Root55             ............MESKSATAA...ASIPAS...TDEANELTGP..VVVAVDVEDEE....
Streptomyces_sp_WM4235             ...........MEQSKSNVAA....HTSAA...VEGTEEITGT..VVVAIDVEDEE....
Streptomyces_sp_MUSC_14            ............MESKSQIAA....TSTP...AEQTEEITGA..VVIAVDVEDEE....
Streptomyces_roseochromogenus      ..........MTSVSQSEGVA.....PAVS...AEQAEEITGT..VVVAVDVEDEE....
Streptomyces_aureocirculatus       ............MVSKSQAAV....STLKP...TQETEELTDT..VVVAVDVEDEE....
Streptomyces_sp_NTK_937            .............MTKSQTTA.AITHAAHA...TQETDELTST..VVVAVDVEDEE....
Streptomyces_sp_DpondAA-D4         .............MSKSQTAA....ATTHA...TQETEELTNT..VVVAVDVEDEE....
```

**Figure S8.** Network 2 overview. (A) Peptide network color-coded by Prodigal score. (B) Representative conserved BGC with PP, YcaO and TfuA genes highlighted. (C) Sequence alignment of all peptides present in network. Network and alignment color-coding is identical to Figure S7.

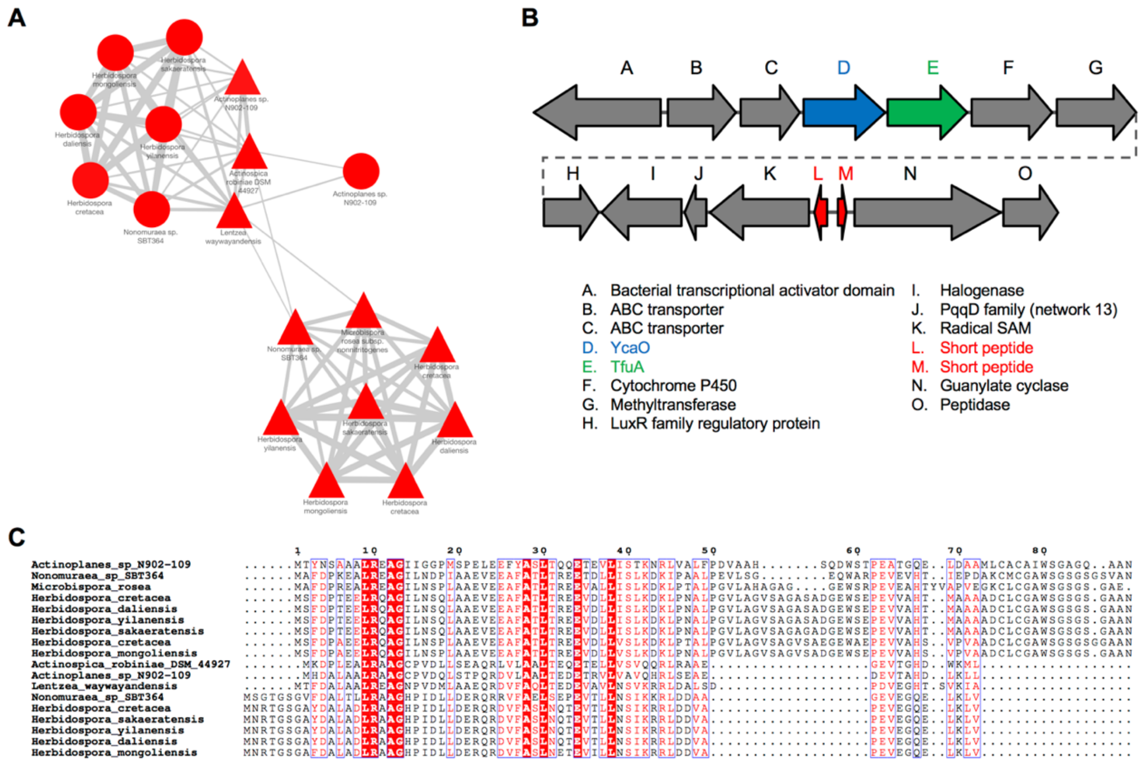


**Figure S9.** Network 3 overview. (A) Peptide network color-coded by Prodigal score. (B) Representative conserved BGC with PP, YcaO and TfuA genes highlighted; some BGCs only have one short peptide in the network. (C) Sequence alignment of all peptides present in network. Network and alignment color-coding is identical to Figure S7.

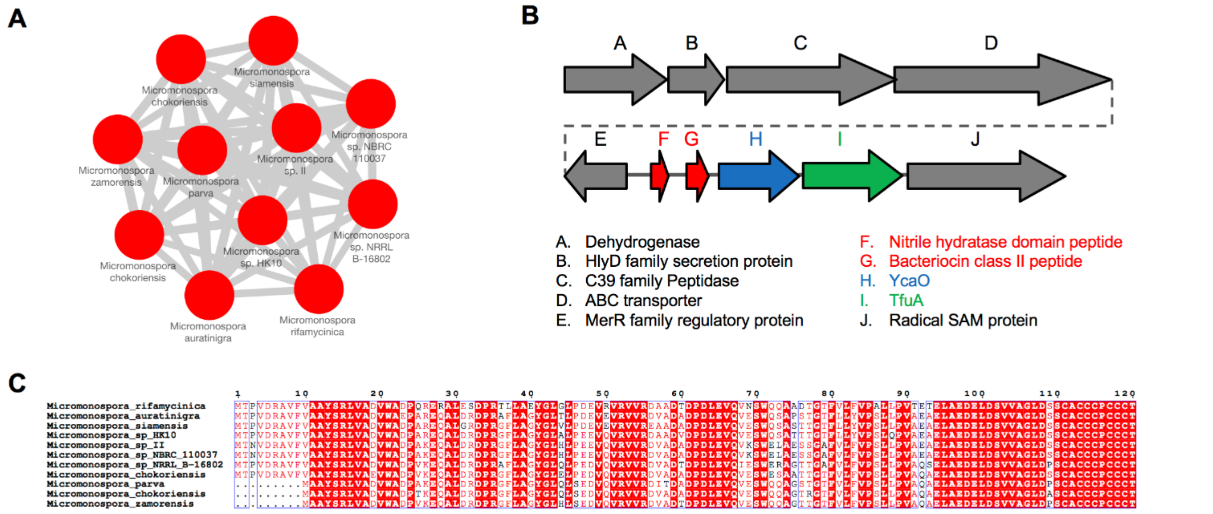


**Figure S10.** Network 7 overview. (A) Peptide network color-coded by Prodigal score. (B) Representative conserved BGC with PP, YcaO and TfuA genes highlighted. The second PP (gene G) was not identified by RiPPER as the peptide is over 120 AA, but MultiGeneBlast indicates homology to the PPs in this network. (C) Sequence alignment of all peptides present in network. Network and alignment color-coding is identical to Figure S7.

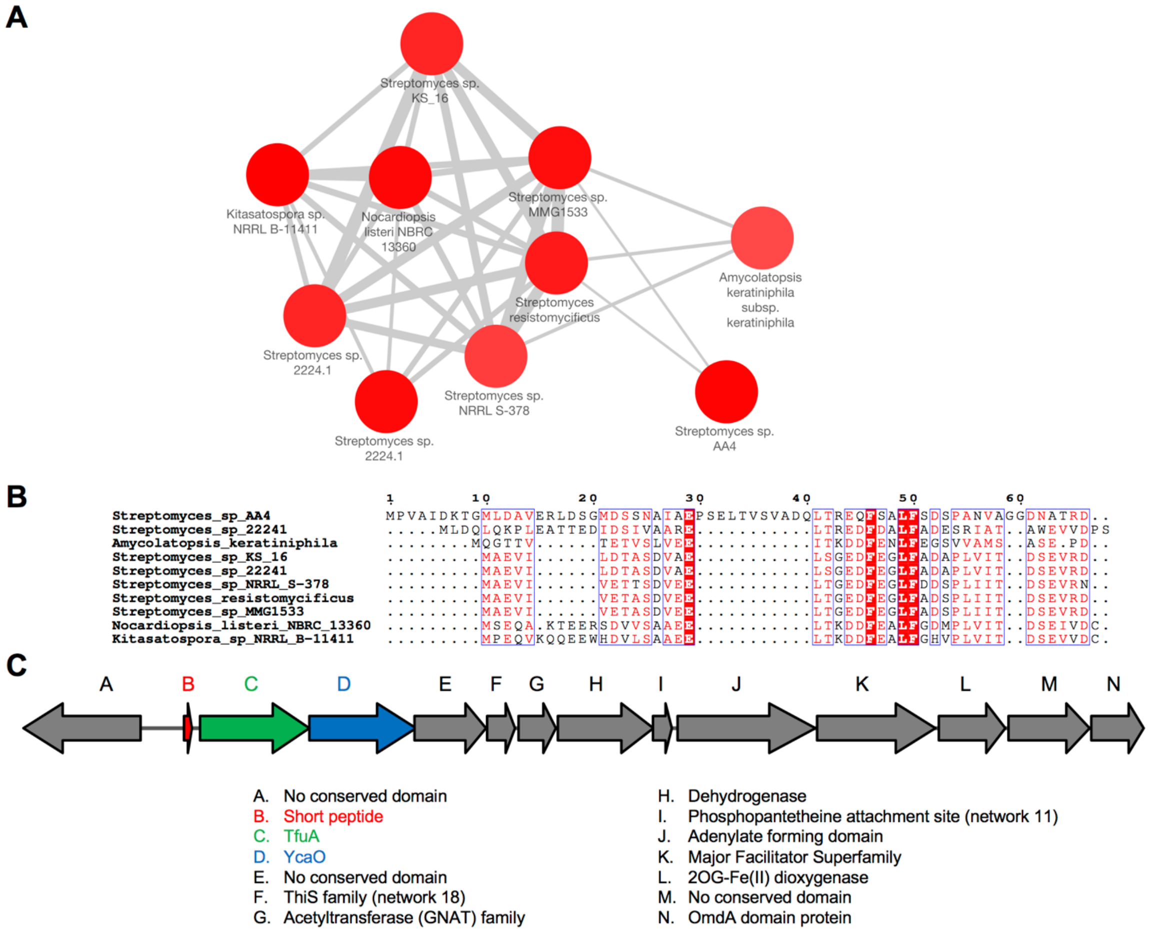


**Figure S11.** Network 9 overview. (A) Peptide network color-coded by Prodigal score. (B) Sequence alignment of all peptides present in network. (C) Representative conserved BGC with PP, YcaO and TfuA genes highlighted. Network and alignment color-coding is identical to Figure S7.

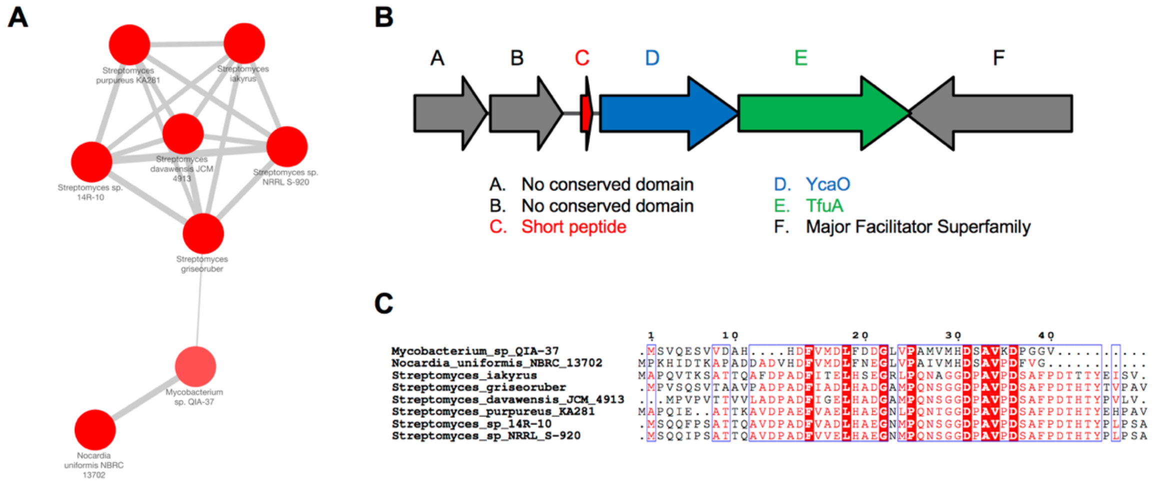


**Figure S12.** Network 12 overview. (A) Peptide network color-coded by Prodigal score. (B) Representative conserved BGC with PP, YcaO and TfuA genes highlighted. (C) Sequence alignment of all peptides present in network. Network and alignment color-coding is identical to Figure S7.

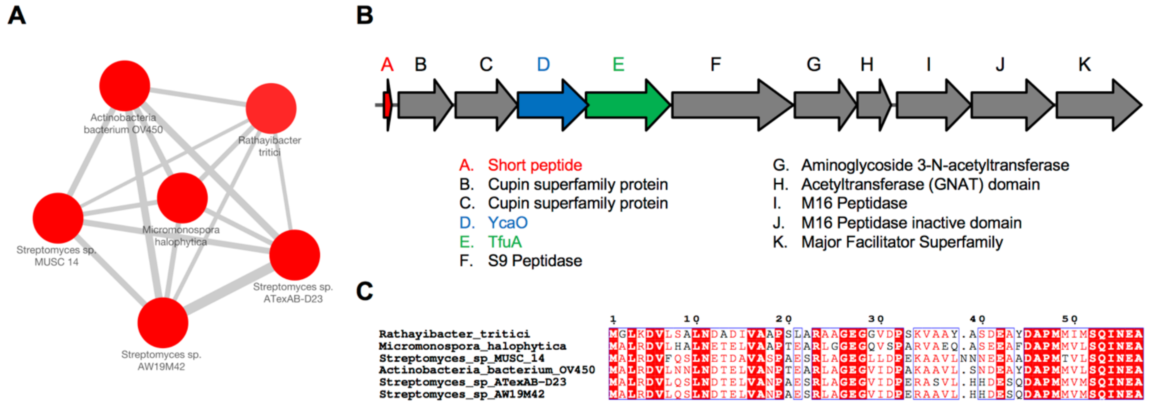


**Figure S13.** Network 15 overview. (A) Peptide network color-coded by Prodigal score. (B) Representative conserved BGC with PP, YcaO and TfuA genes highlighted. (C) Sequence alignment of all peptides present in network. Network and alignment color-coding is identical to Figure S7.

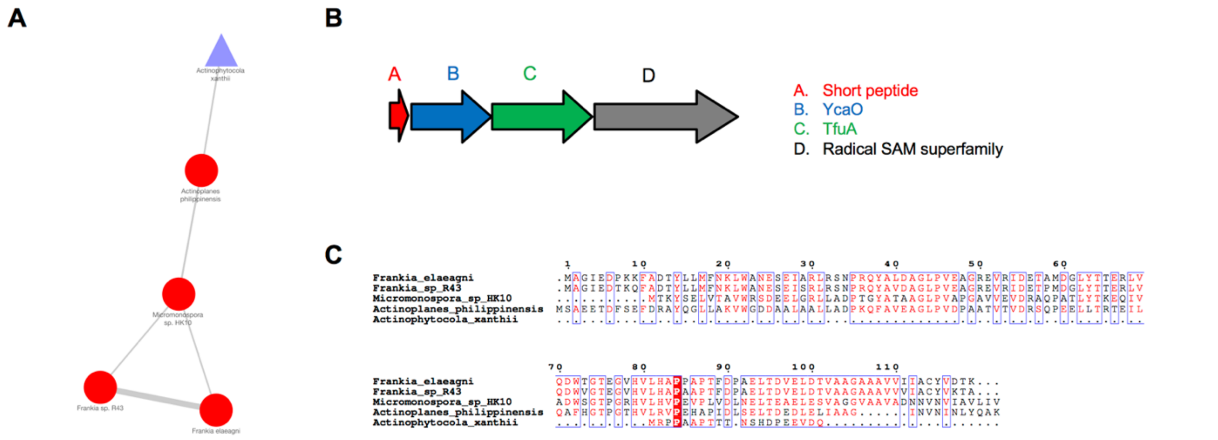


**Figure S14.** Network 20 overview. (A) Peptide network color-coded by Prodigal score. Low score, different orientation, short length and non-canonical position in relation to putative biosynthetic genes indicates that the *Actinophytocola xanthii* peptide is an outlier. (B) Representative conserved BGC with PP, YcaO and TfuA genes highlighted. (C) Sequence alignment of all peptides present in network. Network and alignment color-coding is identical to Figure S7.

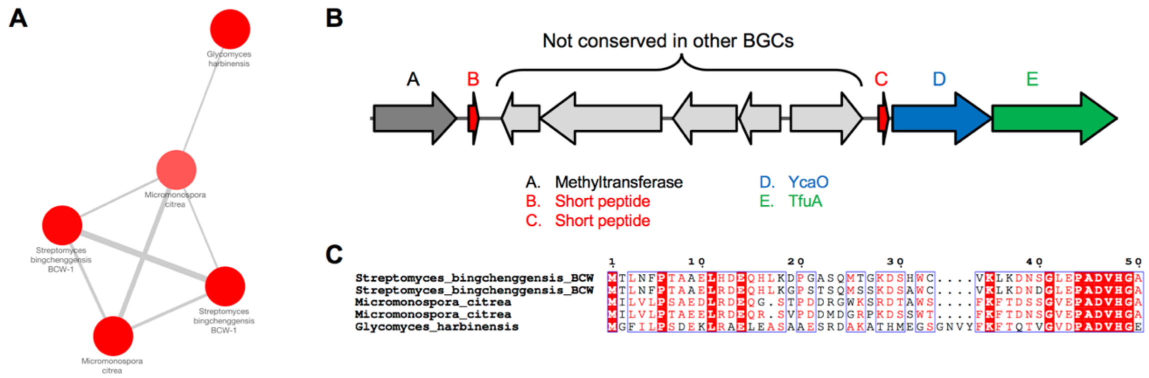


**Figure S15.** Network 21 overview. (A) Peptide network color-coded by Prodigal score. (B) Representative conserved BGC with PP, YcaO and TfuA genes highlighted. Pale grey genes represent an area between short peptides that is not conserved between gene clusters (*Micromonospora citrea* BGC shown). (C) Sequence alignment of all peptides present in network. Network and alignment color-coding is identical to Figure S7.

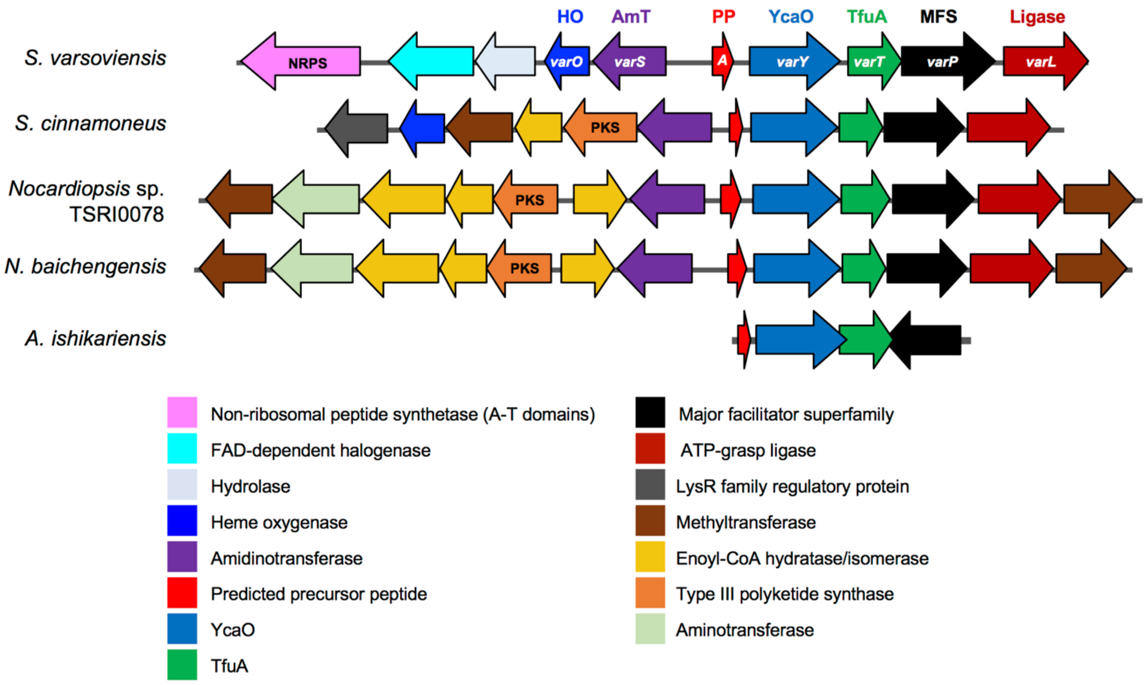


**Figure S16.** Full annotation of the thiovarsolin-like BGCs (Network 22) shown in Fig. 5B.

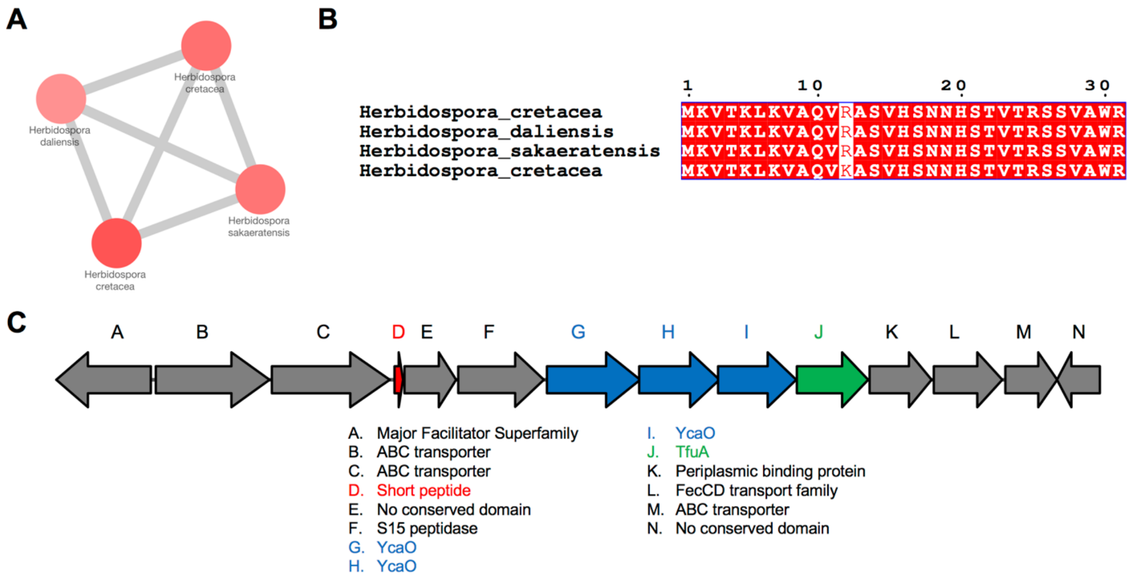


**Figure S17.** Network 24 overview. (A) Peptide network color-coded by Prodigal score. (B) Sequence alignment of all peptides present in network. (C) Representative conserved BGC with PP, YcaO and TfuA genes highlighted. Network and alignment color-coding is identical to Figure S7.

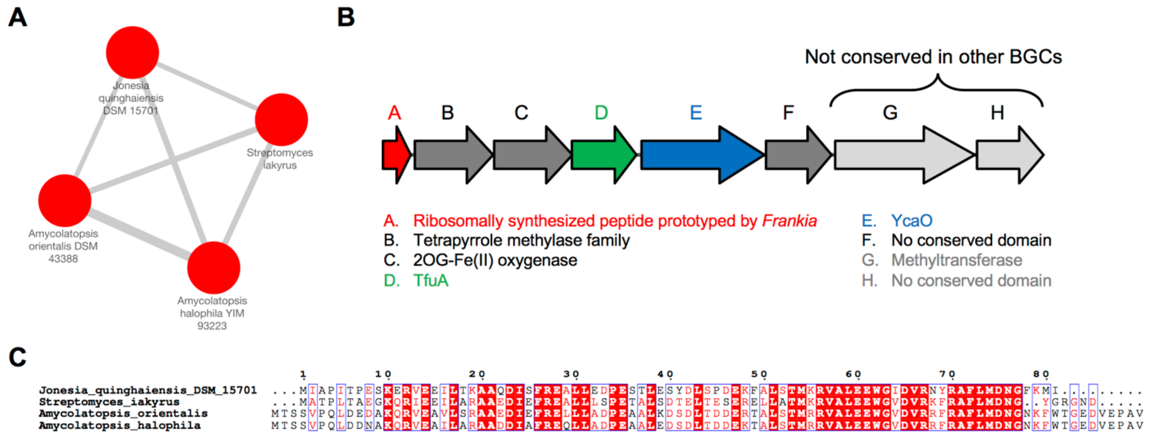


**Figure S18.** Network 27 overview. (A) Peptide network color-coded by Prodigal score. (B) Representative conserved BGC with PP, YcaO and TfuA genes highlighted. Pale grey genes represent an area that is not conserved between gene clusters (the genes shown are found in the *Amycolatopsis* BGCs, but different genes are in the other BGCs). (C) Sequence alignment of all peptides present in network. Network and alignment color-coding is identical to Figure S7.

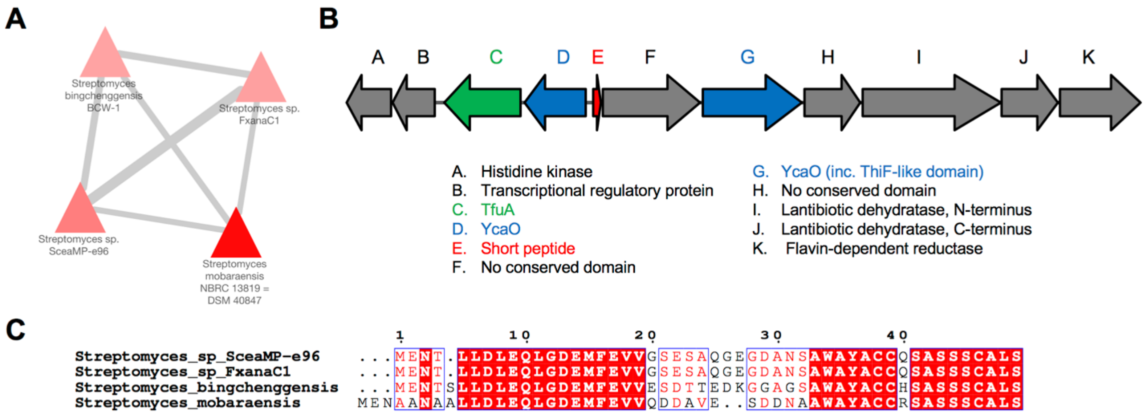


**Figure S19.** Network 28 overview. (A) Peptide network color-coded by Prodigal score. (B) Representative conserved BGC with PP, YcaO and TfuA genes highlighted. (C) Sequence alignment of all peptides present in network. Network and alignment color-coding is identical to Figure S7.

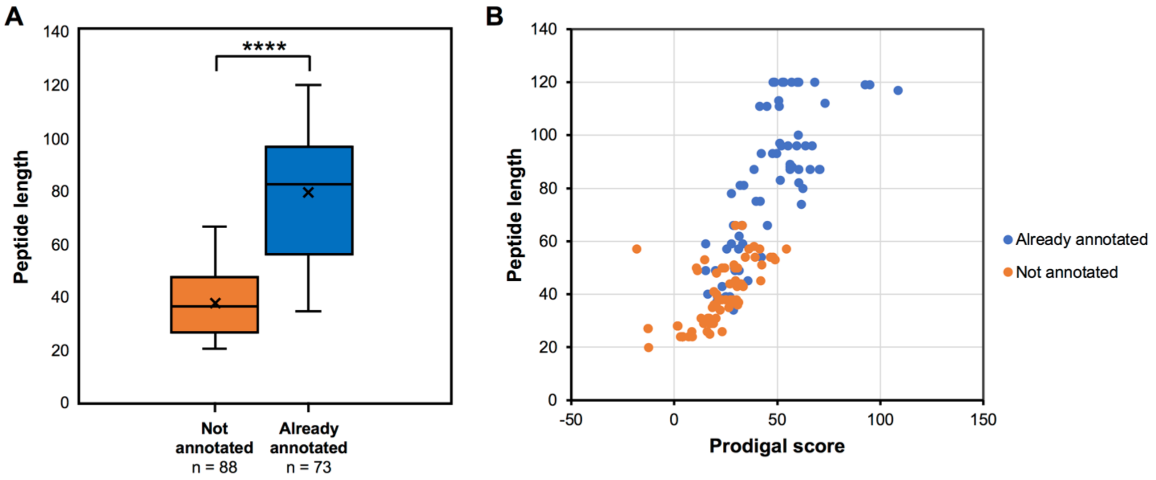


**Figure S20.** Analysis of whether peptides-coding ORFs detected by RiPPER were originally annotated. (A) Comparison of the size distributions of predicted precursor peptides from Networks 1, 2, 3, 5, 7, 9, 12, 15, 20, 21, 22, 24, 26, 27 and 28 that were either previously annotated or not annotated. **** = *p*-value <0.0001 (B) Plot of the peptide length versus the Prodigal score of these peptides.

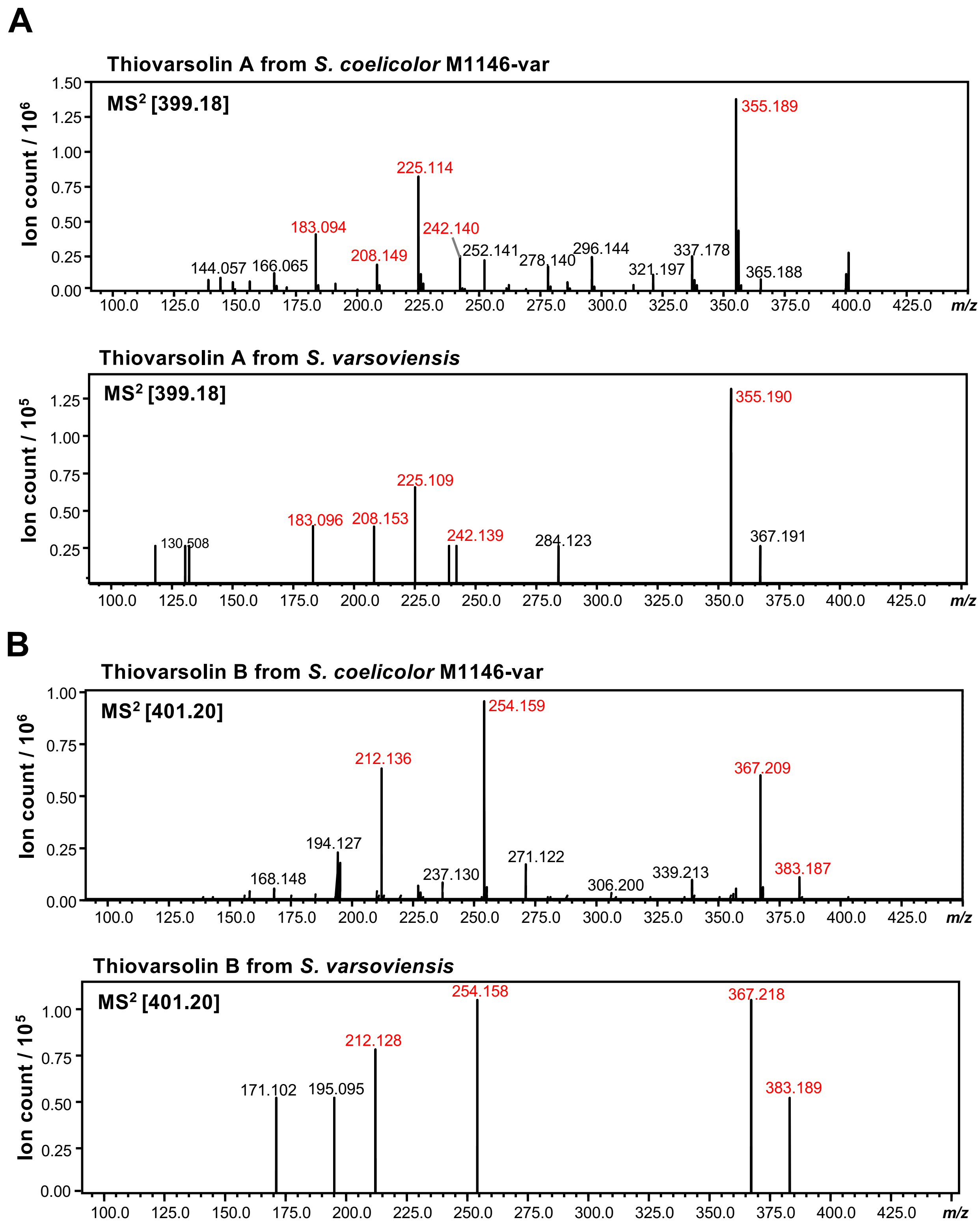


**Figure S21.** Comparsion of $MS^2$ data for thiovarsolins A (panel A) and B (panel B) produced by *S. coelicolor* M1146-var and wild type *S. varsoviensis*. Fragments found in both strains are colored red.

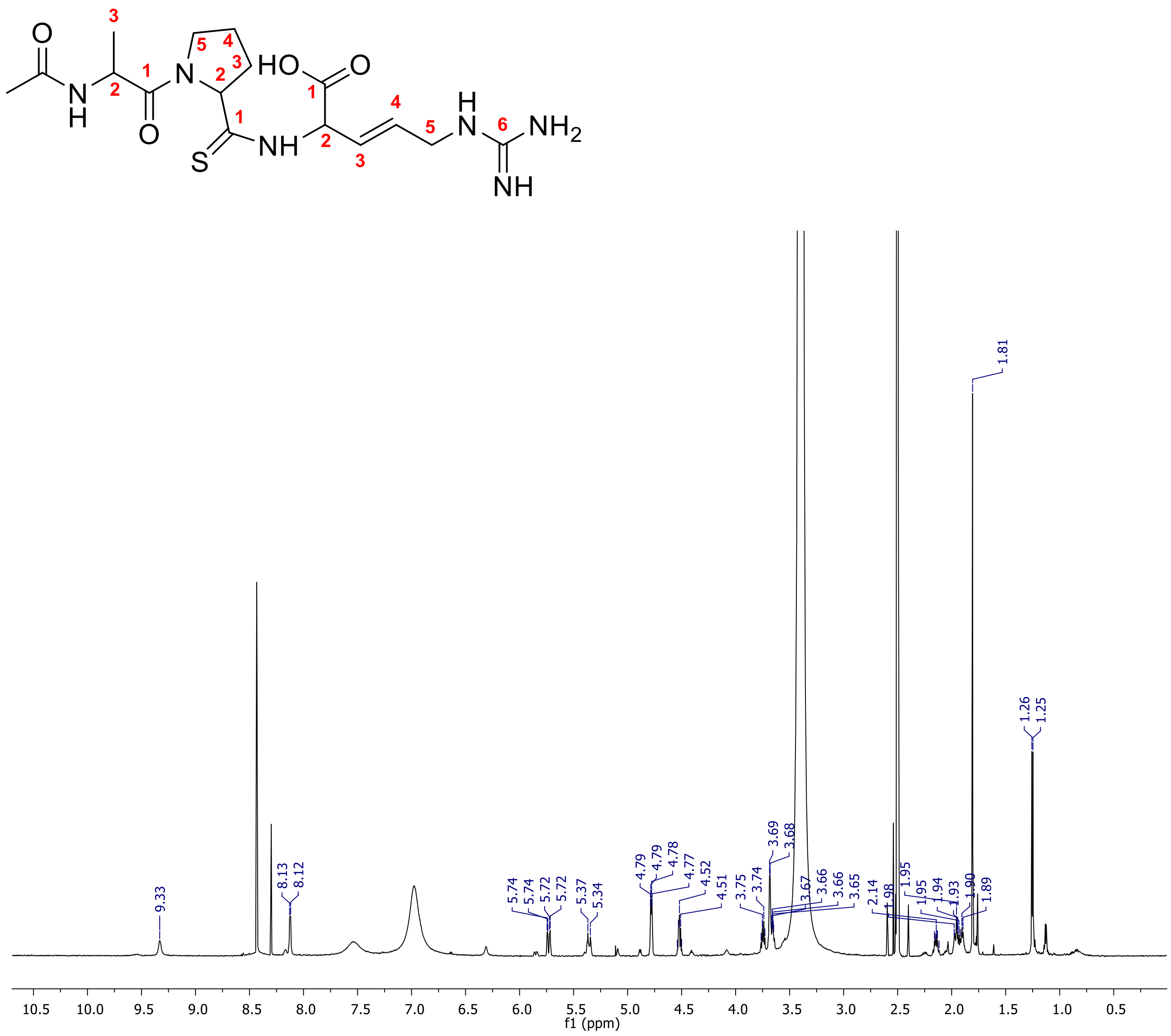


**Figure S22.** 700 MHz $^1$H NMR spectrum of thiovarsolin A in DMSO-$d_6$. Carbon numbering used in Table S6 is also shown.

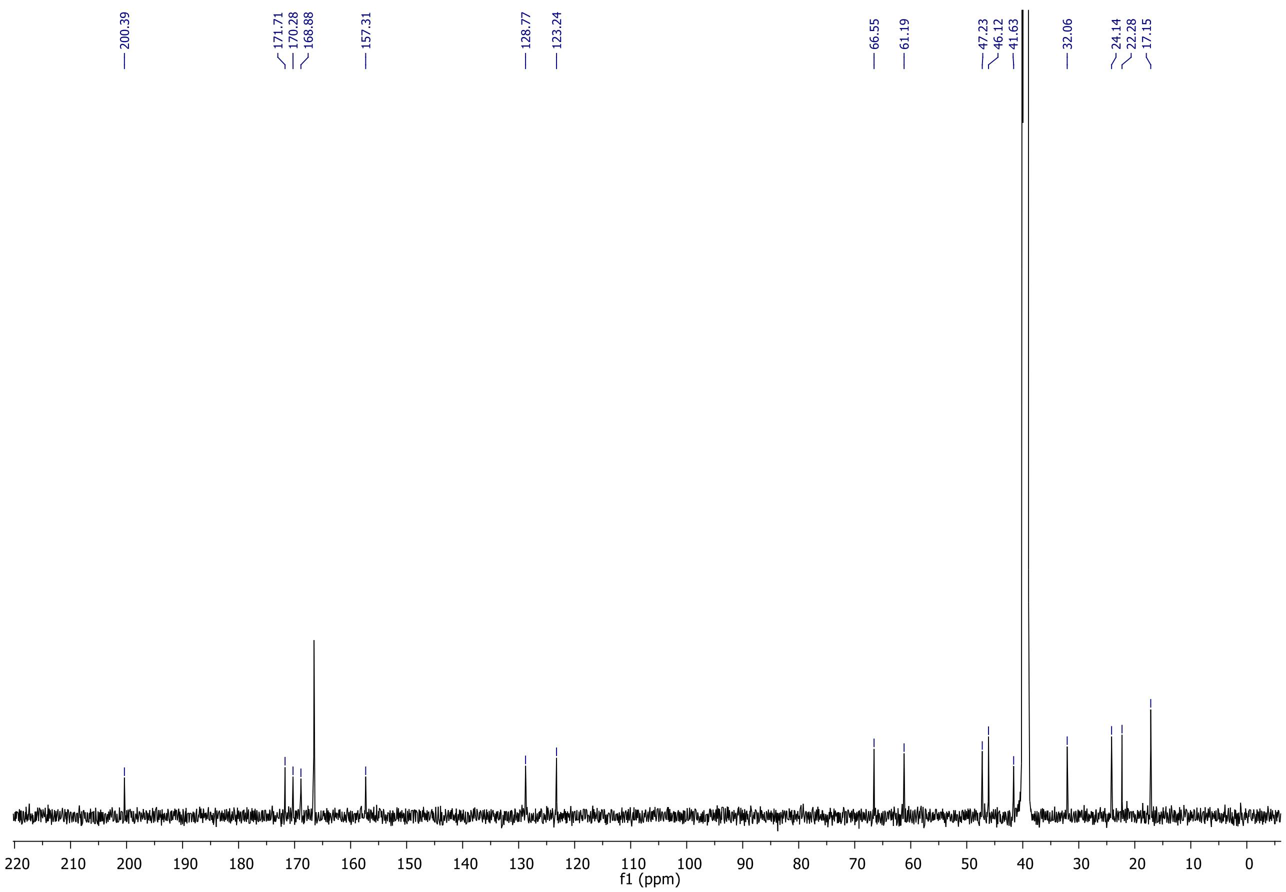


**Figure S23.** 175 MHz $^{13}C$ NMR spectrum of thiovarsolin A in DMSO-$d_6$.

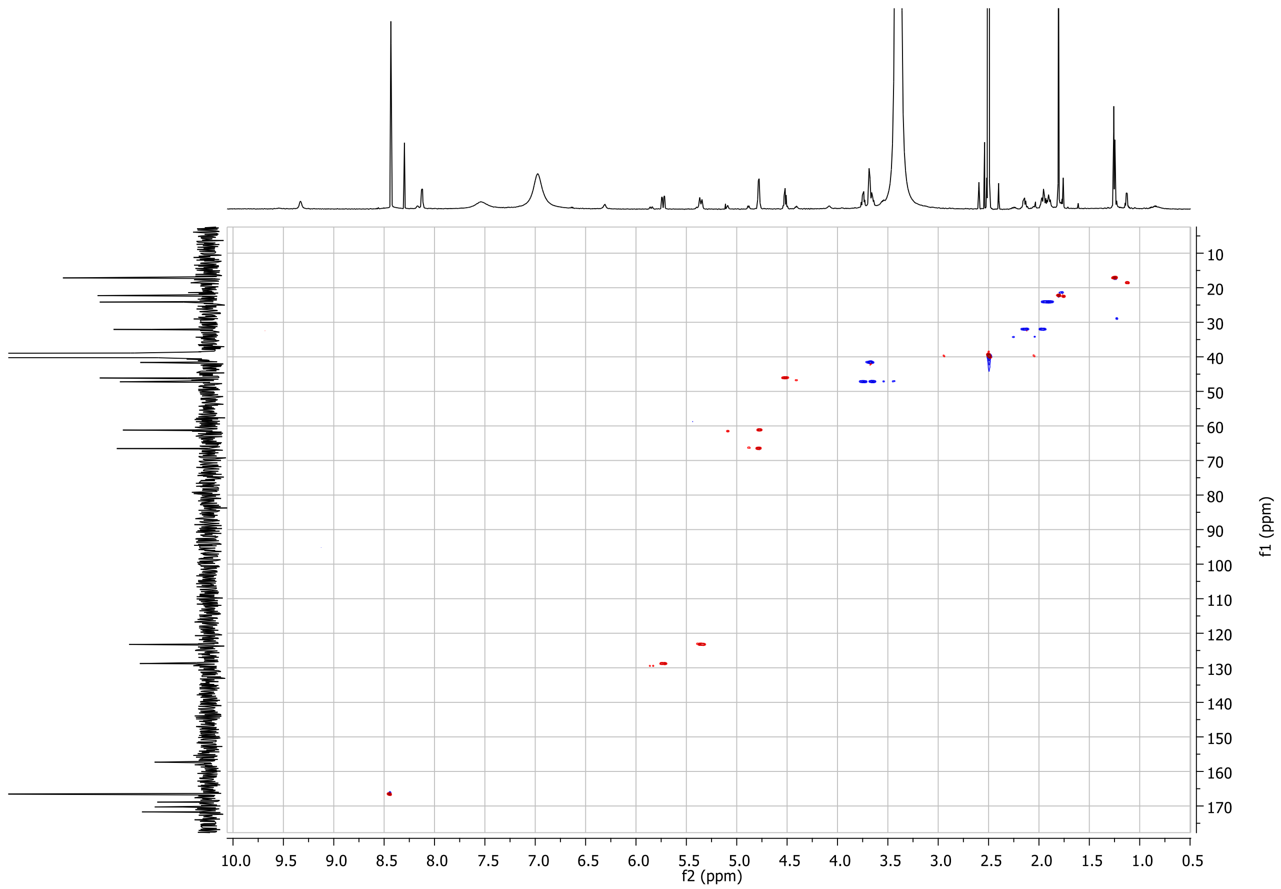


**Figure S24.** 700 MHz $^{1}H$-$^{13}C$ HSQC NMR spectrum of thiovarsolin A in DMSO-$d_6$.

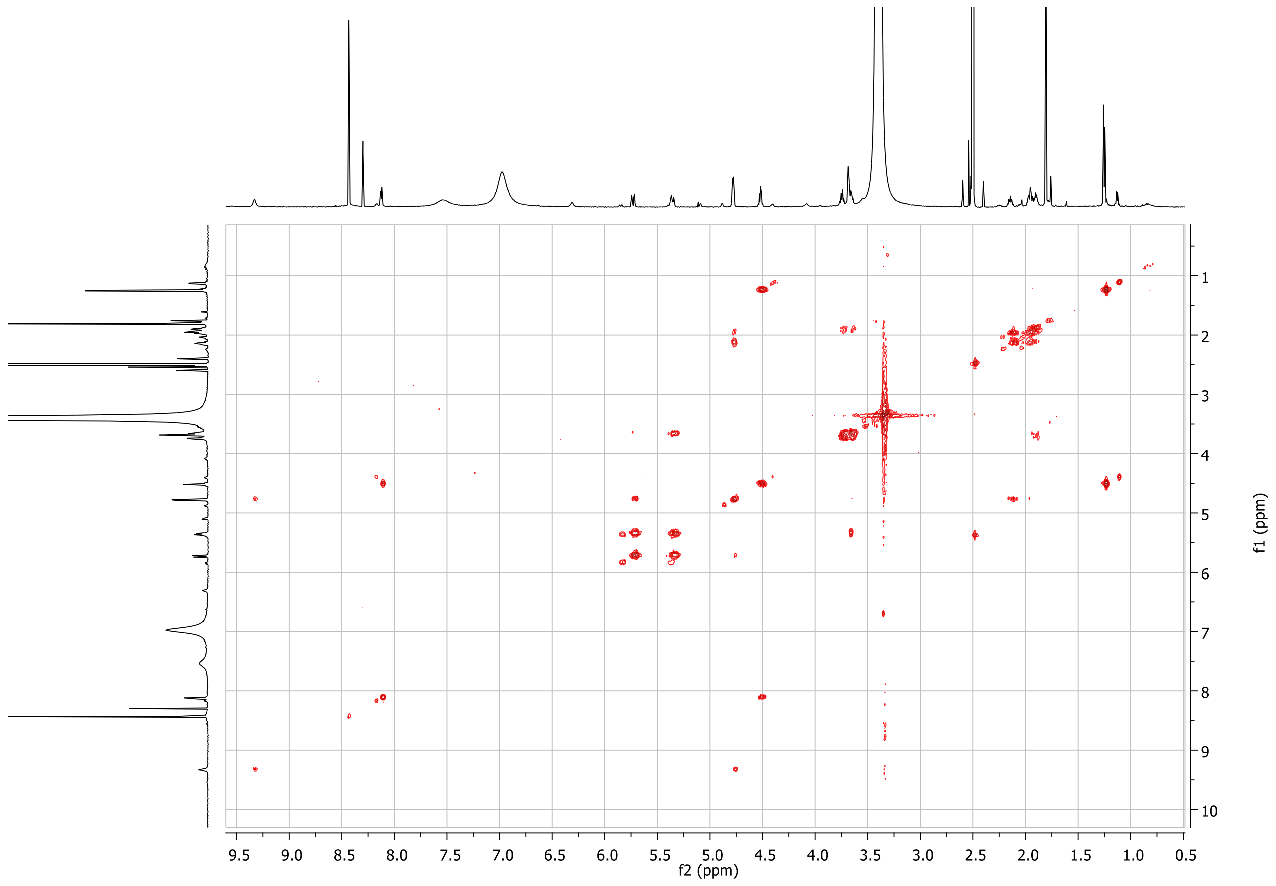


**Figure S25.** 700 MHz $^{1}$H-$^{1}$H COSY NMR spectrum of thiovarsolin A in DMSO-$d_6$.

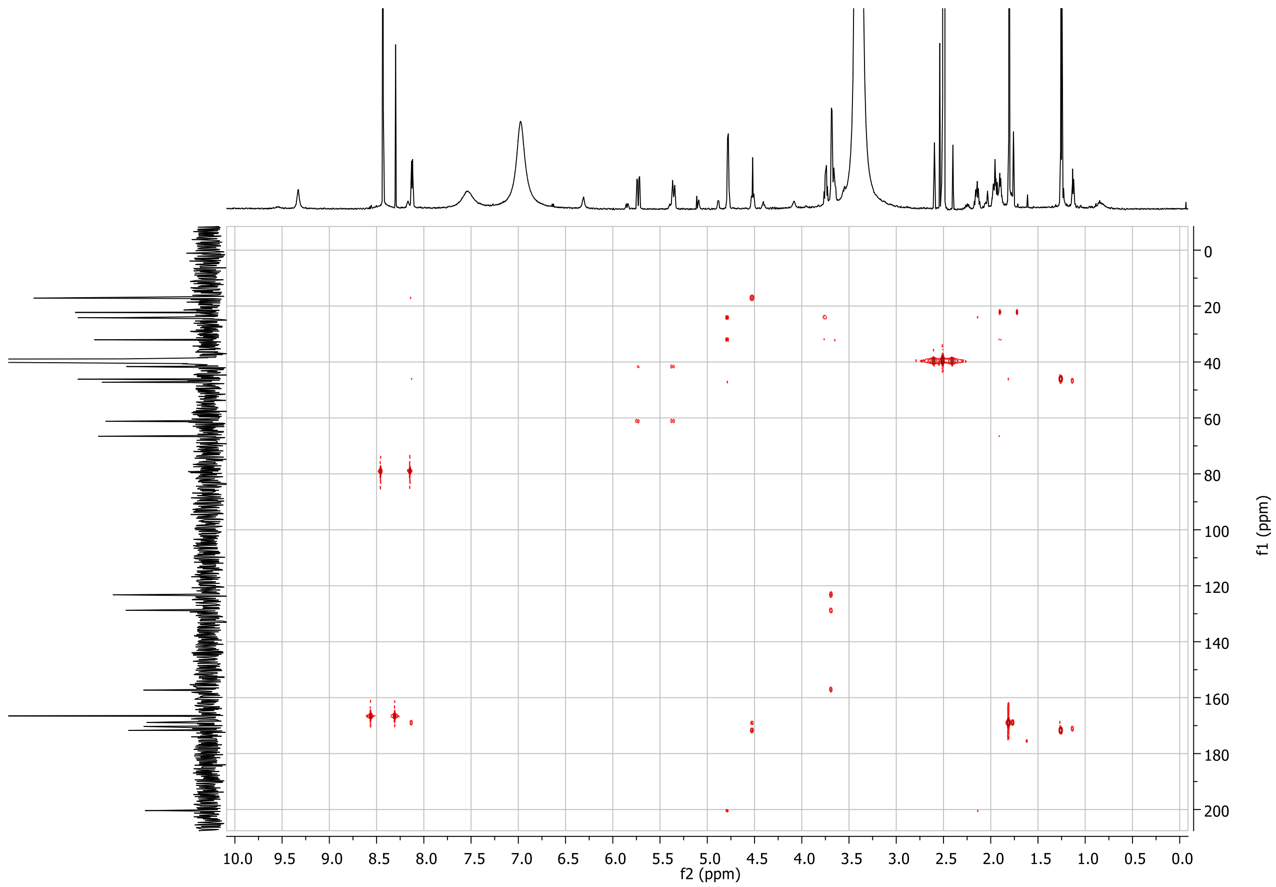


**Figure S26.** 700 MHz $^{1}H$-$^{13}C$ HMBC NMR spectrum of thiovarsolin A in DMSO-$d_6$.

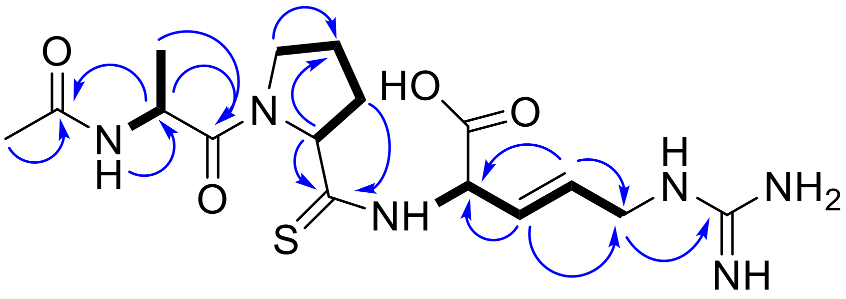


**Figure S27.** $^1$H-$^1$H (COSY, bold lines) and $^1$H-$^{13}$C (HMBC, blue arrows) correlations in thiovarsolin A.

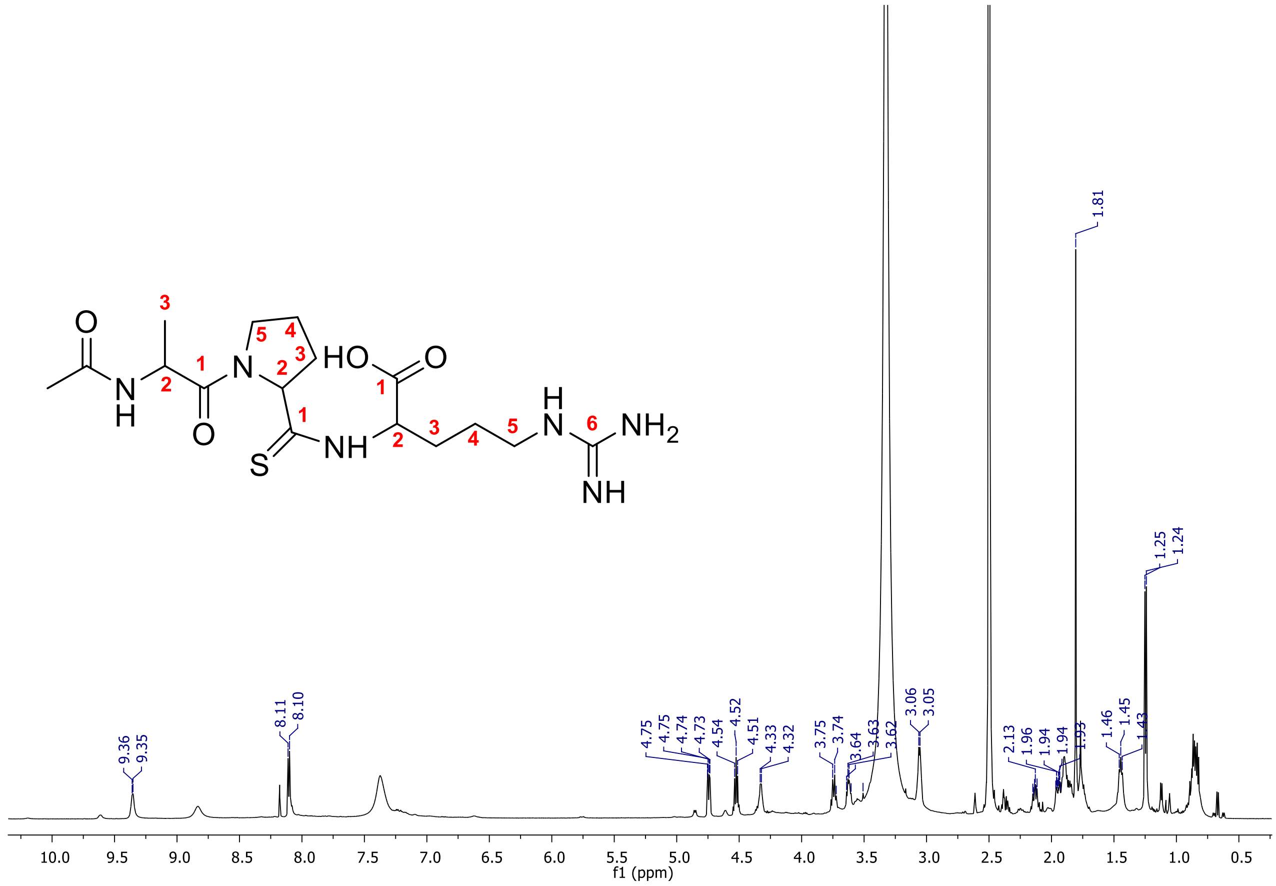


**Figure S28**. 600 MHz $^1$H NMR spectrum of thiovarsolin B in DMSO-$d_6$.

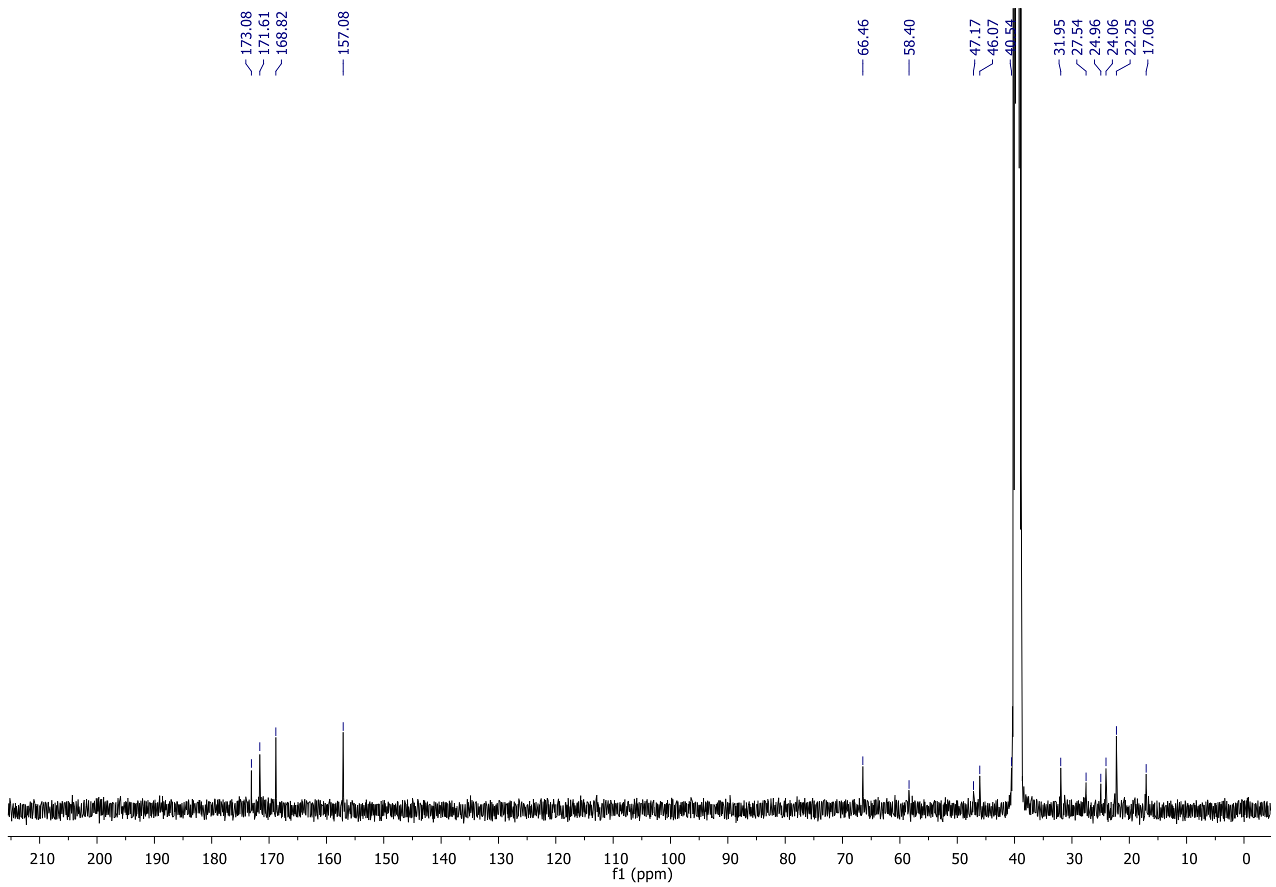


**Figure S29.** 100 MHz $^{13}C$ NMR spectrum of thiovarsolin B in DMSO-$d_6$. The thioamide carbonyl signal is not visible here but is clearly seen in the HMBC spectrum (Figure S32).

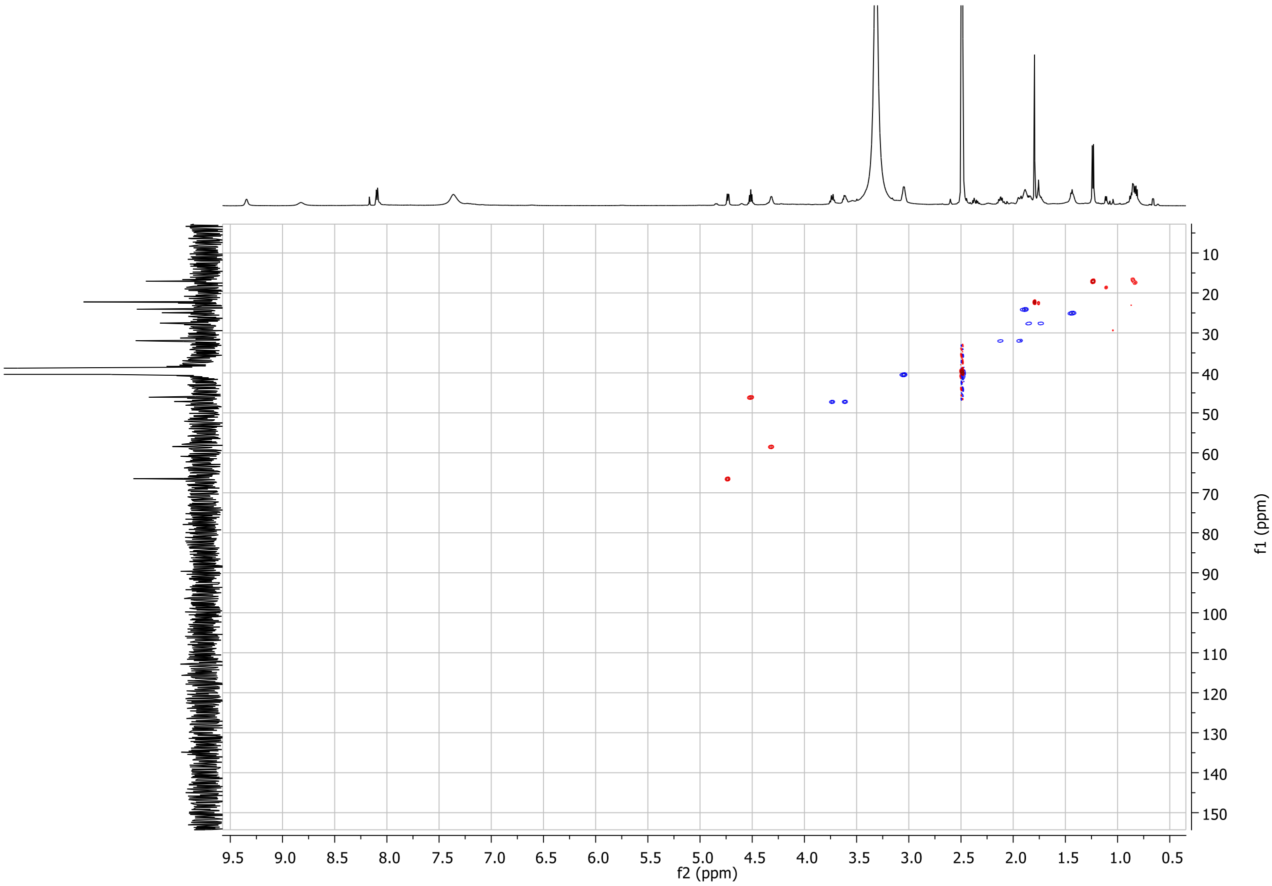


**Figure S30.** 600 MHz $^{1}H$-$^{13}C$ HSQC NMR spectrum of thiovarsolin B in DMSO-$d_6$.

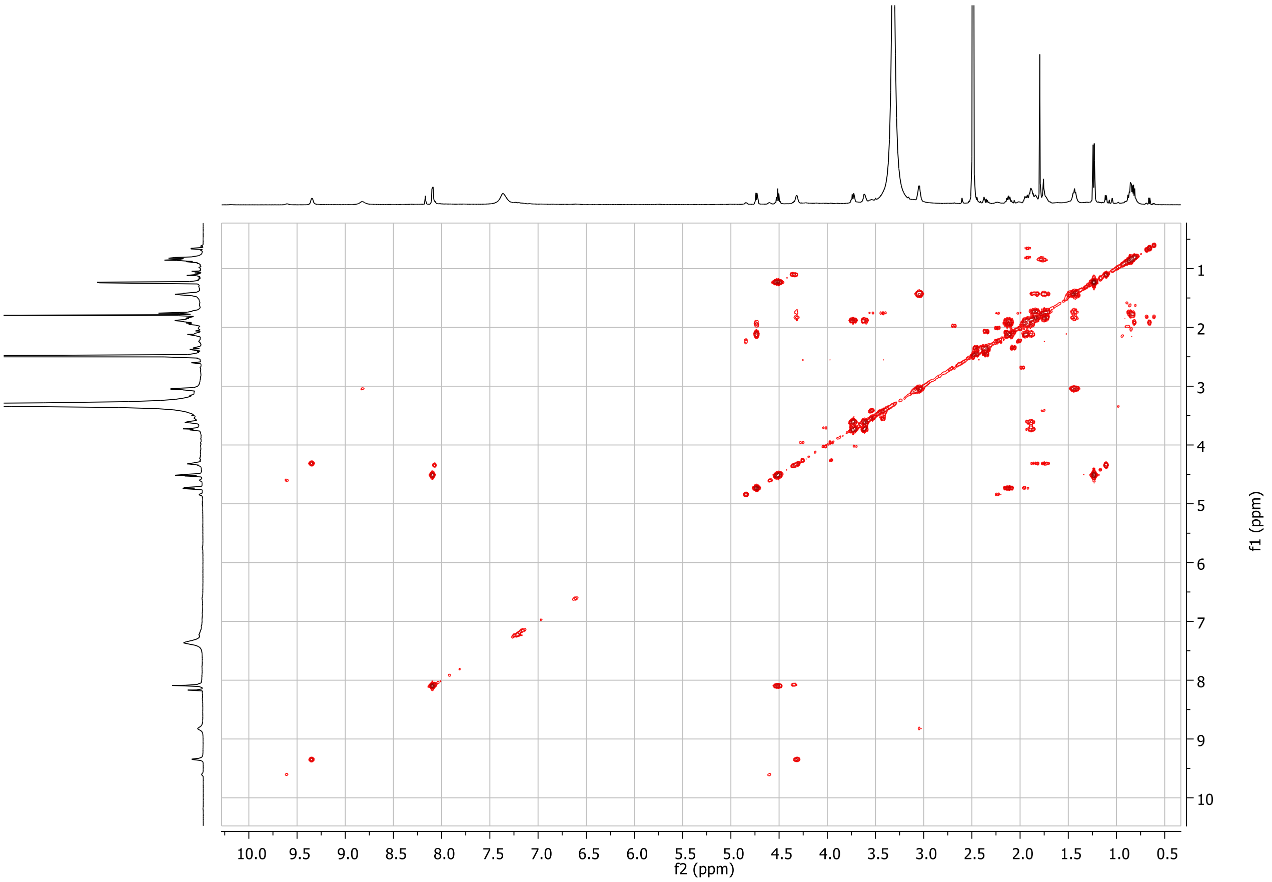


**Figure S31.** 600 MHz $^1H$-$^1H$ COSY NMR spectrum of thiovarsolin B in DMSO-$d_6$.

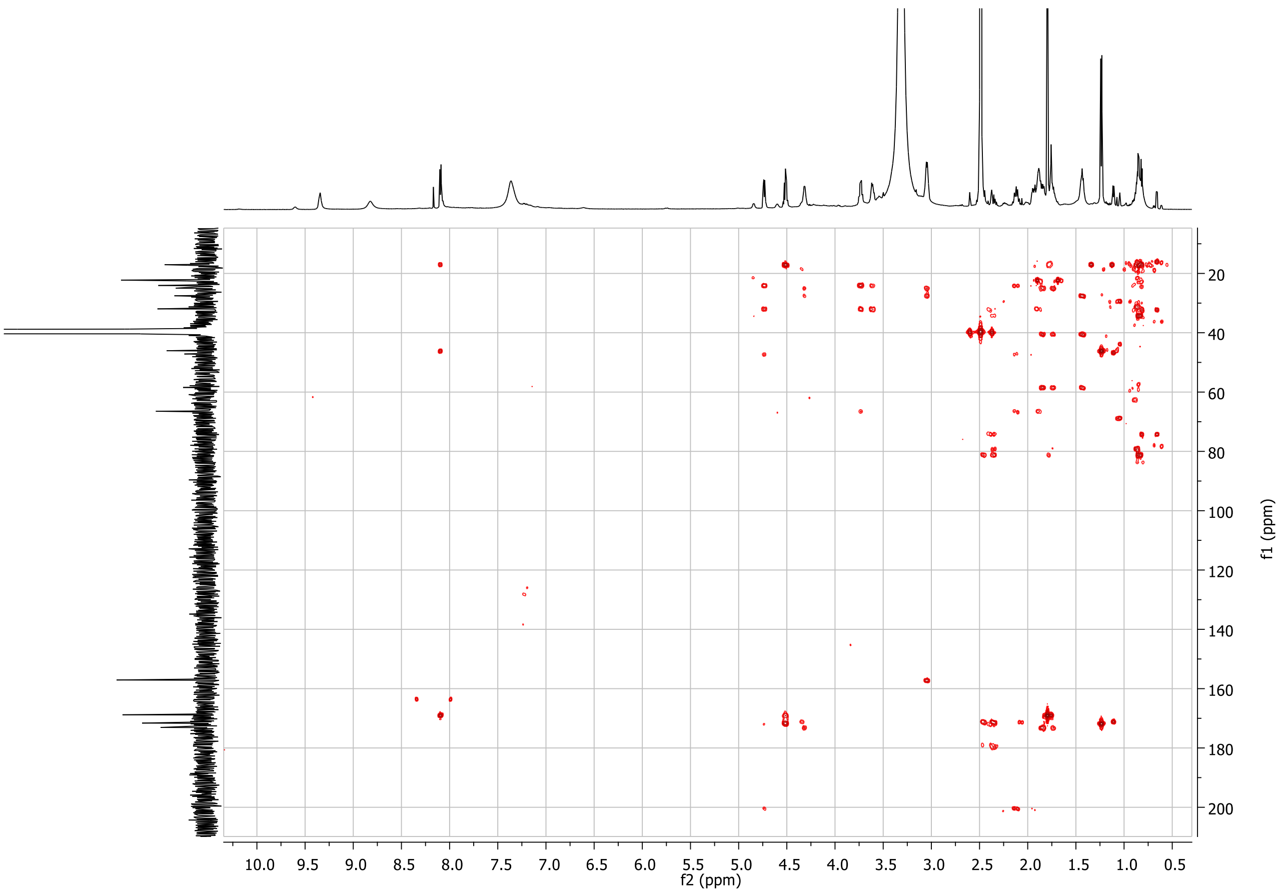


**Figure S32.** 600 MHz $^{1}H$-$^{13}C$ HMBC NMR spectrum of thiovarsolin B in DMSO-$d_6$.

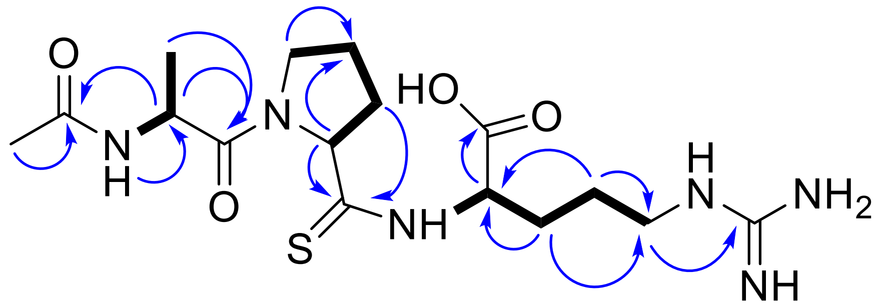


**Figure S33.** $^1$H-$^1$H (COSY, bold lines) and $^1$H-$^{13}$C (HMBC, blue arrows) correlations in thiovarsolin B.

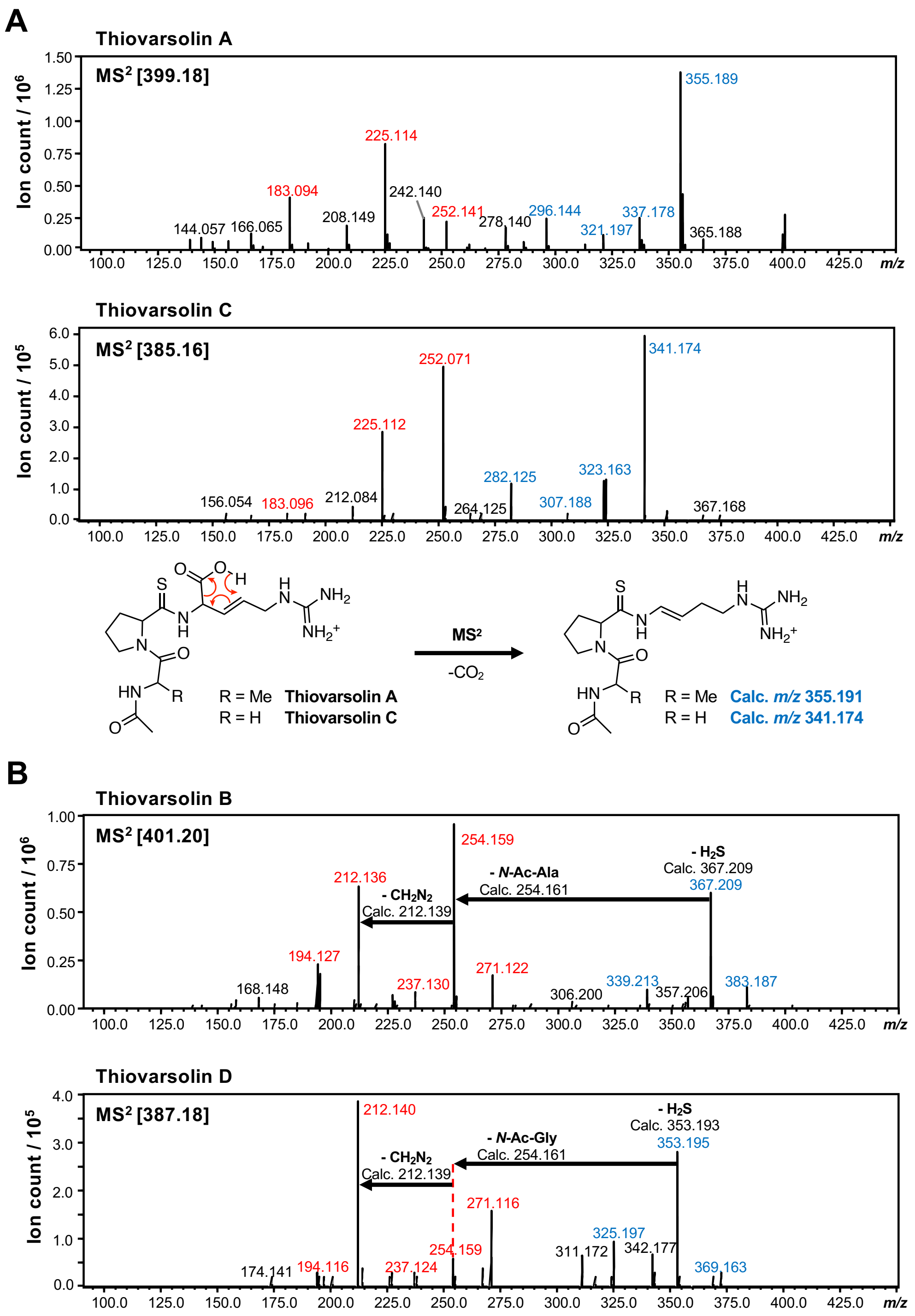


**Figure S34.** MS$^2$ analysis of thiovarsolins produced by *S. coelicolor* M1146-var. (A) Comparison of MS$^2$ spectra of thiovarsolins A and C. The fragmentation data is consistent with a decarboxylation that may be assisted by the double bond on the arginine, as illustrated. (B) Comparison of MS$^2$ spectra of thiovarsolins B and D. In the absence of the Arg double bond, the loss of $H_2S$ dominates, which is characteristic of thioamides.[22] Loss of the acetylated alanine or glycine provides common fragments ($-CH_2N_2$ corresponds to fragmentation on the Arg side chain and is equivalent to an arginine to ornithine fragmentation).

```
VarA      1 MRKDSDLRLDDLAHVNPEQLQGFLEERSTAAVHGATDAYFSGPRAQTVRG     50
            ||||||||||||||||||||||||||||||||||||||||||||||||||
VarA*     1 MRKDSDLRLDDLAHVNPEQLQGFLEERSTAAVHGATDAYFSGPRAQTVRG     50

VarA     51 ATDAHFSAPRAQTVRGATDAHFSAPRAQTVRGATDAHFSAPRAQTVRGAT    100
            ||||:||.||||||||||||..||.||||||||||||:||.|||||||||
VarA*    51 ATDAYFSGPRAQTVRGATDGTFSGPRAQTVRGATDAYFSGPRAQTVRGAT    100

VarA    101 DSYFSAPRAQNVR    113
            |||||.|||||||
VarA*   101 DSYFSGPRAQNVR    113
```

**Figure S35.** Sequence alignment of wild type VarA and the mutated VarA*, where each repeat has been modified to resemble the native GPR-containing repeat. The APR and GPR regions that the thiovarsolins derive from are highlighted in bold and are underlined, and the natural GPR-containing repeat is highlighted in red. Grey shading highlights a two-AA mutation that was a consequence of the approach required to construct such a repetitive gene.

### SUPPLEMENTARY TABLES

**Table S1.** Summary of top 30 peptide networks identified for *tfuA*-containing BGCs in Actinobacteria.

| Network | Consensus Pfam domain | Same strand | Notes | Likely precursor peptide? |
|---|---|---|---|---|
| **1** | None | Yes | *Mycobacteria* | **Possible** |
| **2** | None | Yes | *Streptomyces* | **Yes** |
| **3** | None | Mix | Mainly *Herbidospora*; some BGCs have two peptides in this network | **Possible** |
| **4** | None | Yes | *Streptomyces*; same BGCs as some network 2 peptides; not present in all BGCs with similar tailoring genes. | **No** |
| **5** | None | Yes | Mixed *Actinobacteria*; TLM network | **Yes** |
| **6** | None | No | *Mycobacteria*; same BGCs as some network 1 peptides; low peptide score | **No** |
| **7** | Nitrile hydratase | Yes | *Micromonospora*; Possible second related PP in each cluster not picked as over 120 AA | **Yes** |
| **8** | Nucleoporin C-terminal | No | *Mycobacteria*; same BGCs as some network 1 peptides | **No** |
| **9** | None | Yes | Mixed *Actinobacteria* | **Yes** |
| **10** | DUF732 | No | *Mycobacteria*; same BGCs as some network 1 peptides | **Possible** |
| **11** | Ppant attachment site | Yes | Mixed *Actinobacteria*; same BGCs as some network 9 peptides | **No** |
| **12** | None | Yes | Mainly *Streptomyces* | **Yes** |
| **13** | PqqD | No | Mainly *Herbidospora*; same BGCs as some network 3 peptides | **No** |
| **14** | None | No | Mainly *Micromonospora*; very low peptide scores; same BGCs as some network 7 peptides | **No** |
| **15** | None | Yes | Mixed *Actinobacteria* | **Yes** |
| **16** | Helix-turn-helix | No | *Herbidospora*; same BGCs as some network 3 peptides | **No** |
| **17** | LuxR family | No | Mixed *Actinobacteria*; same BGCs as some network 3 peptides | **No** |
| **18** | ThiS family | Yes | *Streptomyces*; same BGCs as some network 9 peptides | **No** |
| **19** | DUF397 | No | Mixed *Actinobacteria* | **No** |
| **20** | None | Yes | Mixed *Actinobacteria*; *Actinophytocola* entry likely an outlier (low score, limited homology) | **Yes** |

| | | | | |
|---|---|---|---|---|
| **21** | None | Yes | Mixed *Actinobacteria*; Some BGCs have two peptides in this network | **Yes** |
| **22** | None | Yes | Mixed *Actinobacteria*; thiovarsolin network | **Yes** |
| **23** | DUF3017 | Yes | *Mycobacteria*; same BGCs as some network 1 peptides | **Possible** |
| **24** | None | Yes | *Herbidospora*; three YcaO and one TfuA in each BGC | **Yes** |
| **25** | Trm112p | No | *Mycobacterium gordonae*; same BGCs as some network 1 peptides | **No** |
| **26** | None | No | Two *Amycolatopsis* species; one strain has three almost identical peptides; unusual gene position in relation to *ycaO-tfuA* genes | **Possible** |
| **27** | Ribosomally synthesized Frankia peptide | Yes | Mixed *Actinobacteria* | **Yes** |
| **28** | None | No | *Streptomyces*; peptides before series of RiPP biosynthesis genes on opposite strand (including an additional YcaO protein) | **Yes** |
| **29** | DUF2733 | No | *Micromonospora*; very low peptide scores; same BGCs as some network 7 peptides | **No** |
| **30** | None | No | *Mycobacterium tuberculosis*; very low peptide scores; same BGCs as some network 1 peptides | **No** |

**Table S2.** ORFs and associated predicted protein functions in the pTARvar insert.

| ORF | Accession | Length (bp) | Predicted function | Conserved domain | Mutant (PCR targeting primers) | Effect on thiovarsolin production |
|---|---|---|---|---|---|---|
| 1 | WP_030877102.1 | 2097 | Peptidase | PF02897 | ΔLeft_arm1 (RD16 and RD17) | No |
| 2 | WP_030877105.1 | 1101 | AraC regulator / Cupin 6 | PF12833/ PF12852 | ΔLeft_arm1 | No |
| 3 | WP_078644544.1 | 1278 | Major Facilitator Superfamily | PF07690 | ΔLeft_arm1 | No |
| 4 | WP_078644578.1 | 1194 | Methyltransferase | PF01135 | ΔLeft_arm1 | No |
| 5 | WP_030877113.1 | 567 | Nucleoside deaminase | PF00383 | ΔLeft_arm1 | No |
| 6 | WP_030877115.1 | 1962 | NRPS (A/T domains) | PF00501/ PF00550 | ΔLeft_arm2 (RD18 and RD19) | No |
| 7 | WP_030877117.1 | 1407 | Halogenase | PF04820 | ΔLeft_arm2 | No |
| 8 | WP_030877120.1 | 972 | Thioesterase | PF00975 | ΔLeft_arm2 | No |
| 9 (*varO*) | WP_030877123.1 | 690 | Heme oxygenase | PF14518 | Δ*varO* (RD14 and RD15) | **Yes** |
| 10 (*varS*) | WP_030877126.1 | 1134 | Amidinotransferase[a] | Not assigned | Δ*amdT* (RD12 and RD13) | No |
| 11 (*varA*) | WP_030877127.1 | 342 | Precursor Peptide | Not assigned | Δ*varA* (RD1 and RD3) Δ*varA*core (RD2 and RD3) | **Yes** |
| 12 (*varY*) | WP_030877129.1 | 1347 | YcaO-domain protein | PF02624 | Δ*varY* (RD4 and RD5) | **Yes** |
| 13 (*varT*) | WP_063763981.1 | 816 | TfuA-like protein | PF07812 | Δ*varT* (RD6 and RD7) | **Yes** |
| 14 (*varP*) | WP_063763980.1 | 1452 | Major Facilitator Superfamily | PF07690 | Δ*mfs* (RD8 and RD9) | No |
| 15 (*varL*) | WP_030877136.1 | 1362 | ATP-grasp ligase[a] | Not assigned | Δ*atp_gl* | No |

| | | | | | | |
|---|---|---|---|---|---|---|
| | | | | | (RD10 and RD11) | |
| 16 | WP_030877138.1 | 999 | Metallo-dependent hydrolase | PF04909 | ΔRight_arm (RD20 and RD21) | No |
| 17 | WP_030877139.1 | 753 | Short chain Acyl-CoA dehydrogenase | PF13561 | ΔRight_arm | No |
| 18 | WP_030877141.1 | 318 | Hypothetical protein | Not assigned | ΔRight_arm | No |
| 19 | WP_030877143.1 | 1290 | Membrane protein | PF10011 | ΔRight_arm | No |
| 20 | WP_048832106.1 | 1245 | Acyl-CoA transferase | PF02515 | ΔRight_arm | No |
| 21 | WP_078644541.1 | 1653 | Major Facilitator Superfamily | PF07690 | ΔRight_arm | No |
| 22 | WP_030877149.1 | 687 | GntR family regulator | PF00392 | ΔRight_arm | No |
| 23 | WP_030877152.1 | 849 | Formate dehydrogenase associated protein | PF02634 | ΔRight_arm | No |
| 24 | WP_030877153.1 | 474 | MarR family regulator | PF12802 | ΔRight_arm | No |
| 25 | WP_030877155.1 | 933 | NADPH-quinone reductase | PF00107 | ΔRight_arm | No |

[a] Functions predicted by homology to proteins in pheganomycin biosynthesis.[23]

**Table S3.** Oligonucleotides used in this study.

| **Name**[a] | **Sequence**[b] **(5'→3')** |
|---|---|
| TARvar_1 | **TTTGACGCCTCCCATGGTATAAATAGTGGCTCGAGGGGTCGCGGCCTTCTCCGTACCCGCAGCGCTCATCGCCA**CCTCCGCGGGAGTTTAAACCAGGCGATGGC |
| TARvar_2 | **AGCAGCACGTTCCTTATATGTAGCTTTCGACATATGCGAGCAGCGCTCCCGAGGCCAGGGTCAAGGCCCGCCGCAGCCATCGCCT**GGTTTAAACTCCCGCGGAGG |
| CAP03_check-fw | CCGCCTTTTCCTCAATCGCTCTTC |
| CAP03_check-rv | GGACATATCCACGCCCTCCTACATCG |
| TAR_check-fw | GATACAGGATCCGTCATCCCGAGATCCAACGAC |
| TAR_check-rv | GATACAGAATTCCGATCACCCGTACCACGTTGACCG |
| RD1 (RD_*varA*core-fw) | **CAAGGCTTCCTGGAGGAGCGCTCCACCGCCGCCGTCCAC**ATTCCGGGGATCCGTCGACC |
| RD2 (RD_*varA*-fw) | **CGGCCTCCAAACCCGTAAGGAAAGGAAAAGGCCCTCATG**GCAGCTCACGGTAACTGATGC |
| RD3 (RD_*varA*-rv) | **GTGGTCTTTCCGTGGTCTTCCGTGGTCTTCGGGGGCTCA**TGTAGGCTGGAGCTGCTTC |
| RD4 (RD_*varY*-fw) | **CACAGACACGGAGATGGAGACGGAGACAGCGTGAAGATG**GCAGCTCACGGTAACTGATGC |
| RD5 (RD_*varY*-rv) | **GGCTGAGGGGGACTGGGCTGAGGCGAGAGACGCGTGTCA**TGTAGGCTGGAGCTGCTTC |
| RD6 RD_varT-fw: | **GCCTCCGGGCCTGCCTAGCATCTGGGAGCGTTCTCCATG**GCAGCTCACGGTAACTGATGC |
| RD7 RD_varT-rv | **CCGTGGCCGTGGCCCTGCCGCTGTCCGGAGTCTTCTCC**TGTAGGCTGGAGCTGCTTC |
| RD8 RD_msf-fw | **GCACACGCCGCCGGGCCGCTTCGAAAGGGCCCGTTCTTG**GCAGCTCACGGTAACTGATGC |
| RD9 RD_msf-rv | **GGACTCGCGATCGACCACCGCGCCGAAGACGGGCTTCA**TGTAGGCTGGAGCTGCTTC |
| RD10 RD_*atp-gl*-fw | **ATCACCCGCGCCCCCACCCCGAGAAGGGAAGCCACGATG**GCAGCTCACGGTAACTGATGC |
| RD11 RD_*atp-gl*-rv | **GGCGGCGTGGGCCCCAGCCCCACTCGGCAGGGCGGTCA**TGTAGGCTGGAGCTGCTTC |
| RD12 RD_*amt*_fw | **CCCGGTCCTTTCGAAGCCCCAGAGGAGAGCCCATCGATG**GCAGCTCACGGTAACTGATGC |
| RD13 (RD_*amt*-rv) | **CTATGGCGGCCGCGCCCGTGGCCCGCGCCCGGCGGTCA**TGTAGGCTGGAGCTGCTTC |
| RD14 (RD_*varO*-fw) | **GACGAACAACGACGGAAAGCGAGTGGTGGGCGAGCGATG**GCAGCTCACGGTAACTGATGC |

| | |
|---|---|
| RD15 (RD_*varO*-rv) | **CACGGACATGAACATGAACAAGGACAAGGACATGTTCA**TGTAGGCTGGAGCTGCTTC |
| RD16 (RD_left_arm1-fw) | **TGAGGAGCACCGCACGCGGGCGGTCCCGTTCGGCGTCTA**GCAGCTCACGGTAACTGATGC |
| RD17 (RD_left_arm1-rv) | **GAGCGTCCCCGGGCGCCGTCCCACAGGCCCGTCGTCA**TGTAGGCTGGAGCTGCTTC |
| RD18 (RD_left_arm2-fw) | **CGGCGGCCGGCCCCGACGCGGCCTCGGCGGGCCGCGGTG**GCAGCTCACGGTAACTGATGC |
| RD19 (RD_left_arm2-rv) | **TAGACGCCGAACGGGACCGCCCGCGTGCGGTGCTCCTCA**TGTAGGCTGGAGCTGCTTC |
| RD20 (RD_right_arm-fw) | **CCGCCCACCGGCCGCCGGCCGCCGCCCAGGTGATCGTCA**GCAGCTCACGGTAACTGATGC |
| RD21 (RD_right_arm-rv) | **GCACCCACAAGAACCGCACGTCACCTGACTGGGGATCA**TGTAGGCTGGAGCTGCTTC |
| CK1 (*varA*_check-fw) | CGCACAACTCGGCAGAGGCGG |
| CK2 (*varA*_check-rv) | CGGCGGACAGGGACTCGAC |
| CK3 (*varY*_check-fw) | GCCAAGTCGAGTCCCTGTCCG |
| CK4 (*varY*_check-rv) | GGTGAGGCGAGGGGACTGAGAGAG |
| CK5 (*varT*_check-fw) | CCCCTCGCCTCACCGTCCTCTTTC |
| CK6 (*varT*_check-rv) | GGAGACCAGCAGGACGACGC |
| CK7 (*msf*_check-fw) | GCCACGGATTCCGACACGGTC |
| CK8 (*msf*_check-rv) | GCGGGTGATTCGGCTGATTTGG |
| CK9 (*atp-gl*_check-fw) | CCAAATCAGCCGAATCACCCGC |
| CK10 (*atp-gl*_check-rv) | CGGTGGGCGGACGGTCG |
| CK11 (*amT*_check-fw) | CGTTGTTCGTCGCTTTCCCTATGG |
| CK12 (*amT*_check-rv) | CCGTCCCGTCCCCGATGC |
| CK13 *varO*_check-fw) | CCATAGGGAAAGCGACGAACAACG |
| CK14 (*varO*_check-rv) | CATGGACATGAACATGGACATGGC |
| CK15 (left_arm1_check-fw) | GGGAAGCCTACACCGTGCAGTG |
| CK16 (left_arm1_check-rv) | CGCTGTTCGCCACCGTGCTCG |
| CK17 (left_arm2_check-fw) | GCCCGAGAAGCACCCCTACTAGAC |
| CK18 (left_arm2_check-rv) | CCTTGTTCATGTTCATGTCCGTGGC |
| CK19 (right_arm_check-fw) | CGACCGTCCGCCCACCG |
| CK20 (right_arm_check-fw) | GAACCGCACGTCACCTGACTGG |

| | |
|---|---|
| CP1 (*varA*_comp-fw) | GATACACATATGAGGAAAGACTCGGATCTCAGGC |
| CP2 (*varA*_comp-rv) | GATACAAAGCTTCTGTGCCTGTCTGTCTTCGTATTTCTCTG |
| CP3 (*varA*p_comp-fw) | GATACACATATGGGGCTCTCCTCTGGGGCTTCG |
| CP4 (*varA*_comp-rv2) | GATACAAAGCTTCCCGCATCTTCACGCTGTCTCC |
| CP5 (*varY*_comp-fw) | GATACACATATGCGGGCTCACGCCCCC |
| CP6 (*varY*_comp-rv) | GATACAAAGCTTGGAGAACGCTCCCAGATGCTAGG |
| CP7 (*varT*_comp-fw) | GATACACATATGGGACCCGGTGGTCTTCCTCG |
| CP8 (*varT*_comp-rv) | GATACAAAGCTTCGAGCCCGAGCATGGGTACG |
| CP9 (*varO*_comp-fw) | GATACACATATGACGGAGAGTCTGGACCGGG |
| CP10 (*varO*_comp-rv) | GATACAAAGCTTCGTGAGGTCGGTCAGCAGCG |
| AG1 | GTGGACGGCGGCGGTGG |
| AG2 | CCGCGCACCGTCTGGGCCCGCGGGCCCGAGAAGTACGCGTCCGTCGCG<br>CCGTGGACGGCGGCGGTGG |
| AG3 | GATACATCTAGAGGTACCGTCGGTCGCCCCCCGGACGGTCTGCGCCCGC<br>GGCCCGCTGAAGTAGGCGTCGGTCGCCCCGCGCACCGTCTGGGC |
| AG4 | GTCCGCTGAGCCCCCGAAGACC |
| AG5 | GGCGCCACGGACTCCTACTTCTCCGGACCGCGGGCCCAGAACGTCCGCT<br>GAGCCCCCGAAGACC |
| AG6 | GATACAGGTACCTTCAGCGGGCCGCGGGCGCAGACCGTCCGGGGGGCG<br>ACCGACGCCTACTTCAGCGGGCCGCGGGCGCAGACCGTCCGGGGCGCC<br>ACGGACTCCTACTTCTCC |

[a] Primer naming code: RD = primers for PCR targeting; CK = primers for mutant checking; CP = primers for the complementation of mutants; AG = primers for alanine → glycine substitution in *varA*.

[b] Bold letters indicate homologous recombination regions. Underlined letters represent restriction sites.

**Table S4.** Vectors and constructs used in this study.

| Vector / construct | Relevant features | Use | Source or reference |
|---|---|---|---|
| pCAP03 | $kan^R$, *oriT*, ΦC31 int-attP | Capture of the thiovarsolin BGC | 18 |
| pTARvar | $kan^R$, *oriT*, ΦC31 int-attP, 31.7 kb insert from *S. varsoviensis* | Heterologous pathway expression | This work |
| pTARvar Δ*varA*_clean | $kan^R$, *oriT*, ΦC31 int-attP, in-frame deletion of the *varA* core peptide | Functional analysis of *varA* | This work |
| pTARvar Δ*varA* | Same as pTARvar, insertional deletion of *varA* | Functional analysis of *varA* | This work |
| pTARvar Δ*varY* | Same as pTARvar, insertional deletion of *varY* | Functional analysis of *varY* | This work |
| pTARvar Δ*varT* | Same as pTARvar, insertional deletion of *varT* | Functional analysis of *varT* | This work |
| pTARvar Δ*varO* | Same as pTARvar, insertional deletion of *varO* | Functional analysis of *varO* | This work |
| pTARvar Δ*amT* | Same as pTARvar, insertional deletion of *amT* | Functional analysis of *amT* | This work |
| pTARvar Δ*atp_gl* | Same as pTARvar, insertional deletion of *varA* | Functional analysis of *atp_gl* | This work |
| pTARvar Δ*mfs* | Same as pTARvar, insertional deletion of *mfs* | Functional analysis of *mfs* | This work |
| pTARvar Δleft_arm1 | Same as pTARvar, insertional deletion of 5 genes in the left arm | Functional analysis of multiple genes | This work |
| pTARvar Δleft_arm2 | Same as pTARvar, insertional deletion of 3 genes in the left arm | Functional analysis of multiple genes | This work |
| pTARvar Δright_arm | Same as pTARvar, insertional deletion of 10 genes in the left arm | Functional analysis of multiple genes | This work |
| pIJ773 | $amp^R$, $apra^R$ and *oriT* flanked by FRT sites | Template for PCR – targeting cassette | 19 |

| | | | |
|---|---|---|---|
| pIJ773 Δ*oriT* | $amp^R$, $apra^R$ flanked by FRT sites | Template for PCR – targeting cassette | John Innes Centre Collection |
| pIJ10257 | $hyg^R$, oriT, ϕBT1 int-attB, ermEp* | Constitutive expression in *Streptomyces* | 20 |
| pIJ10257_*varA* | Same as pIJ10257, carrying varA | *ΔvarA* complementation | This work |
| pIJ10257_*varA*p | Same as pIJ10257, carrying *varA* and its native promoter | ΔvarA complementation (under *var A* native promoter) | This work |
| pIJ10257_*varY* | Same as pIJ10257, carrying *varY* | *ΔvarY* complementation | This work |
| pIJ10257_*varT* | Same as pIJ10257, carrying *varT* | *ΔvarT* complementation | This work |
| pIJ10257_*varO* | Same as pIJ10257, carrying *varO* | *ΔvarO* complementation | This work |
| pIJ10257_*varA*p*YT* | Same as pIJ10257, carrying am operon comprising *varA*, *varY* and *varT*. | Expression of the minimal thiovarsolins BGCs (under the *varA* native promoter) | This work |
| pGP9 | $apra^R$, oriT, ϕBT1 int-attB, Act-ORFIV / $P_{actI}$ activator / promoter | Constitutive expression in *Streptomyces* | 21 |
| pGP9_*varA**p | pGP9-based, carrying *varA* with a point mutation. | *ΔvarA* complementation with a mutant peptide | This work |

**Table S5.** Accurate mass data for thiovarsolins A-D.

| Compound | $[M+H]^+$ formula | Expected *m/z* | Observed *m/z* | Mass error (ppm) |
|---|---|---|---|---|
| Thiovarsolin A | $C_{16}H_{27}N_6O_4S^+$ | 399.1809 | 399.1818 | +2.25 |
| Thiovarsolin B | $C_{16}H_{29}N_6O_4S^+$ | 401.1966 | 401.1968 | +0.50 |
| Thiovarsolin C | $C_{15}H_{25}N_6O_4S^+$ | 385.1653 | 385.1652 | -0.26 |
| Thiovarsolin D | $C_{15}H_{27}N_6O_4S^+$ | 387.1809 | 387.1808 | -0.26 |

**Table S6.** $^1H$ and $^{13}C$ NMR data for thiovarsolins A and B in DMSO-$d_6$.

| | Thiovarsolin A | | Thiovarsolin B | |
|---|---|---|---|---|
| **Residue**[a] | **$\delta_C$** | **$\delta_H$, mult., *J* in Hz** | **$\delta_C$** | **$\delta_H$, mult., *J* in Hz** |
| **Ala** | | | | |
| 1 | 171.7, C | - | 171.6, C | - |
| 2 | 46.1, CH | 4.52, dd, 7.2, 7.0 | 46.1, CH | 4.52, dq, 7.0, 7.0 |
| 3 | 17.2, $CH_3$ | 1.25, d, 7.0 | 17.1, $CH_3$ | 1.25, d, 7.0 |
| NH | - | 8.12, d, 7.2 | - | 8.11, d, 7.0 |
| CO | 168.9, C | - | 168.8, C | - |
| $CH_3$ | 22.3, $CH_3$ | 1.81, s | 22.3, $CH_3$ | 1.81, s |
| **Pro** | | | | |
| 1 | 200.4, C | - | 200.2, C[b] | - |
| 2 | 66.6, CH | 4.78, dd, 7.9, 4.5 | 66.5, CH | 4.74, dd, 8.6, 3.0 |
| 3 | 32.1, $CH_2$ | 1.95, m<br>2.14, m | 32.0, $CH_2$ | 1.95, ddd, 11.5, 8.6<br>2.13, m |
| 4 | 24.1, $CH_2$ | 1.92, m | 24.1, $CH_2$ | 1.87, m |
| 5 | 47.2, $CH_2$ | 3.66, m<br>3.75, dd, 14.6, 7.4 | 47.2, $CH_2$ | 3.62, dd, 14.0, 8.0<br>3.74, dd, 14.0, 8.6 |
| **Arg'** | | | | |
| 1 | 170.3, C | - | 173.1, C | - |
| 2 | 61.2, CH | 4.78, dd, 7.9, 4.5 | 58.4, CH | 4.33, dd, 6.4, 5.7 |
| 3 | 128.8, CH | 5.73, dd, 15.6, 7.9 | 27.5, $CH_2$ | 1.74, m<br>1.84, m |
| 4 | 123.4, CH | 5.36, dd, 15.6, 4.5 | 25.0, $CH_2$ | 1.42, m |
| 5 | 41.6, $CH_2$ | 3.68, d, 4.5 | 40.5, $CH_2$ | 3.08, m |
| 6 | 157.3, C | - | 157.1, C | - |
| NH | - | 9.33, br s | | 9.35, d, 6.4 |

[a] Carbon numbering is shown in Figures S22 and S28.

[b] Assigned based on HMBC correlations (Figure S32).
