## Supplementary figures and images for "Uncovering the unexplored diversity of thioamidated ribosomal peptides in Actinobacteria using the RiPPER genome mining tool"

### Supplementary file 6

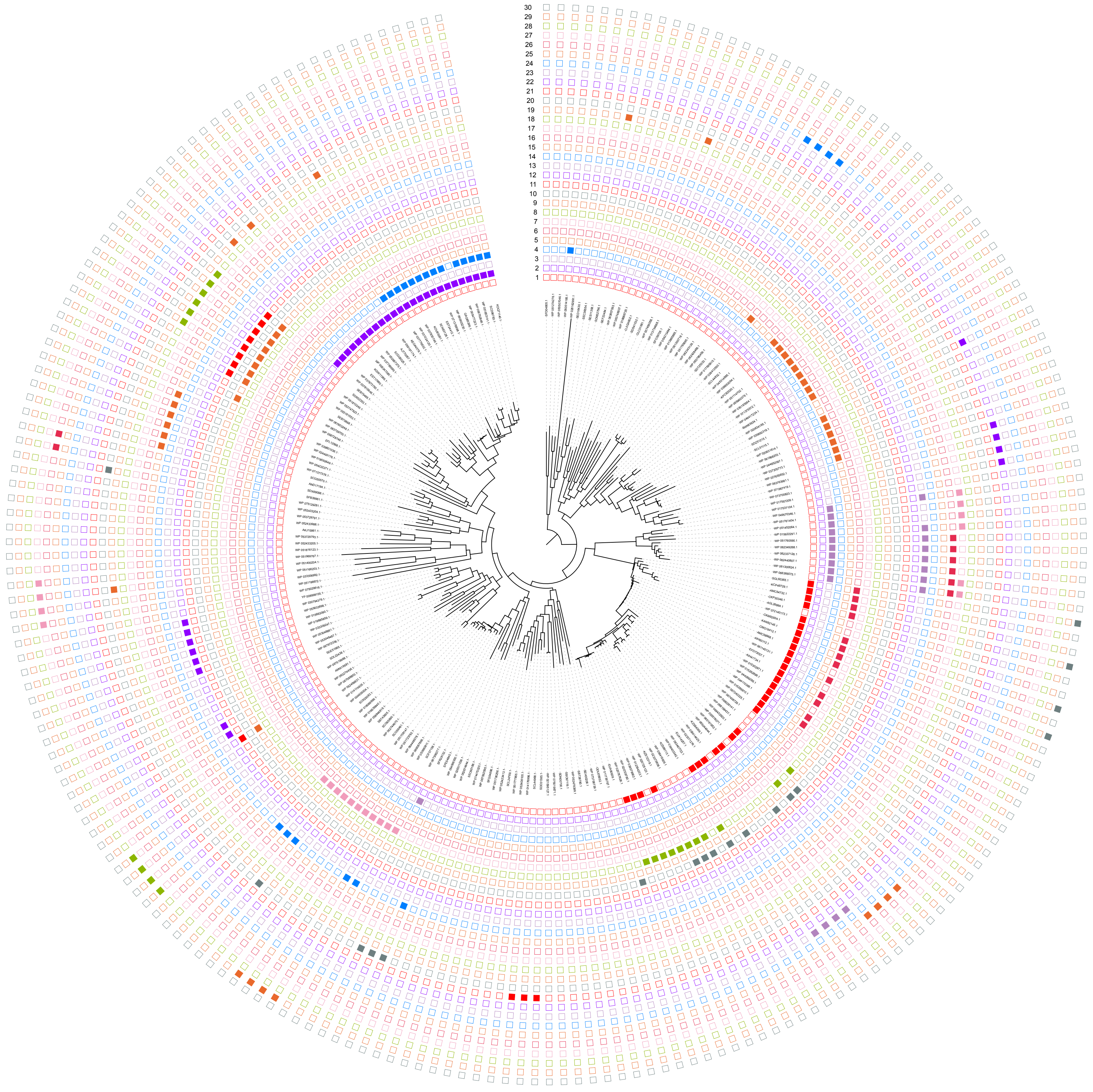
